# A molecular toolbox for ADP-ribosyl binding proteins

**DOI:** 10.1101/2021.05.31.445082

**Authors:** Sven T. Sowa, Albert Galera-Prat, Sarah Wazir, Heli I. Alanen, Mirko M. Maksimainen, Lari Lehtiö

**Author notes:** These authors contributed equally (Sven T. Sowa, Albert Galera-Prat).

## Abstract

Proteins interacting with ADP-ribosyl groups are often involved in disease-related pathways or in viral infections, which makes them attractive targets for the development of inhibitors. Our goal was to develop a robust and accessible assay technology that is suitable for high-throughput screening and applicable to a wide range of proteins acting as either hydrolysing or non-hydrolysing binders of mono- and poly-ADP-ribosyl groups. As a foundation of our work, we developed a C-terminal protein fusion tag based on a Gi protein alpha subunit peptide (GAP), which allows for site-specific introduction of cysteine-linked mono- and poly-ADP-ribosyl groups as well as chemical ADP-ribosyl analogs. By fusion of the GAP-tag and ADP-ribosyl binders to fluorescent proteins, we were able to generate robust FRET signals and the interaction with 22 previously described ADP-ribosyl-binders was confirmed. To demonstrate the applicability of this binding assay for high-throughput screening, we utilized it to screen for inhibitors of the SARS-CoV-2 nsp3 macrodomain and identified the drug suramin as a moderate yet unspecific inhibitor of this protein. To complement the binding technology, we prepared high-affinity ADP-ribosyl binders fused to a nanoluciferase, which enabled simple blot-based detection of mono- and poly-ADP-ribosylated proteins. These tools can be expressed recombinantly in *E. coli* using commonly available agents and will help to investigate ADP-ribosylation systems and aid in drug discovery.

## Introduction

ADP-ribosylation is a post-translational modification involved in the regulation of many diverse processes in the cell. Despite its physiological importance, the intricate interplay of ADP-ribose transfer, detection and removal and its connection to specific pathways is not well-understood at the molecular level (Gupte et al., 2017; O’Sullivan et al., 2019; Palazzo and Ahel, 2018). ADP-ribosyl-transferases (“writers”) such as PARP family members catalyse the transfer of mono- or poly-ADP-ribosyl groups to target proteins. These ADP-ribosyl groups can then be recognized by ADP-ribosyl-binding proteins (“readers”), which often serve to recruit further proteins. ADP-ribosyl groups may also be removed by hydrolysing proteins (“erasers”), reversing the action of writers and thereby impacting respective signalling processes.

The human genome encodes many different readers and erasers of ADP-ribosyl groups. Macrodomains represent a large class of ADP-ribose binders known in humans. Many of them are encoded as part of other multidomain proteins such as ADP-ribosyltransferases (PARP9, PARP14, PARP15) or histones (macroH2A variants). Other macrodomains such as MDO1, MDO2, PARG or TARG1 possess hydrolytic activity and are integral actors in ADP-ribose signalling pathways (Feijs et al., 2013; Rack et al., 2016, 2020a). Viruses such as coronaviruses or togaviruses are known to harbour macrodomains that can remove ADP-ribose from proteins inside the host cell. These viral macrodomains are implied to weaken the host virus defence mechanism by interfering with the host ADP-ribosylation signalling machinery and have been shown to be necessary for virus replication and pathogenesis (Abraham et al., 2018; Fehr et al., 2018; Leung et al., 2018; McPherson et al., 2017). Viruses with these macrodomains include Chikungunya virus, MERS-CoV (camel flu) and SARS-CoV-2 (COVID-19). In addition to macrodomains, ADP-ribosyl glycohydrolases of another human enzyme family, ARH1 and ARH3, have been reported to hydrolyse ADP-ribose linked especially to arginine and serine residues, respectively (Abplanalp et al., 2017; Rack et al., 2018, 2020a). Several other domain types exist that are primarily associated with binding of poly-ADP-ribosyl groups (PAR). Examples of these are the PAR-binding zinc finger (PBZ) domain of APLF (Li et al., 2010; Rulten et al., 2008), WWE domain in multiple ADP-ribosyltransferases and E3 ubiquitin ligases (Aravind, 2001; Wang et al., 2012) and PAR-binding phosphate pocket in the BRCT1 domain of XRCC1 (Breslin et al., 2015; Kim et al., 2015).

While development of inhibitors for ADP-ribosyltransferases has been investigated for multiple decades mainly in the context of cancer therapeutics, the development of inhibitors against binders or hydrolysers of ADP-ribose has only gained momentum in recent years (Palazzo and Ahel, 2018). Inhibitors of ADP-ribose binding and hydrolysing proteins would be valuable tools that could help to decipher the complex ADP-ribosyl signalling machinery inside the cell. These inhibitors might also display therapeutic potential as recently demonstrated with novel PARG inhibitors that can impair cancer cell survival (Houl et al., 2019). Furthermore, the inhibition of viral macrodomains to combat diseases caused by these viruses was suggested (Fehr et al., 2018; Rack et al., 2020b) and is currently being explored extensively for the SARS-CoV-2 nsp3 macrodomain (Shimizu et al., 2020).

We sought to set up a system that enables easy development of binding assays suited for a wide variety of ADP-ribosyl-binders and -hydrolases alike, could further be used for inhibitor screening in a high-throughput setting, and would be simple and easily accessible. Many previous binding assays could only be applied to either ADP-ribosyl readers (Ekblad et al., 2018) or erasers (Haikarainen et al., 2018; Wazir et al., 2021) or are not suitable for high-throughput setups like mass-spectrometry and immunoblot based methods (Haikarainen and Lehtiö, 2016; Hirsch et al., 2014). Many assays also rely on expensive reagents as used in AlphaScreen or TR-FRET technologies or use custom synthesized reagents. Voorneveld et al. developed a method to synthesize peptides containing mono-ADP-ribosyl (MAR) groups in a selected serine, threonine or cysteine residue (Voorneveld et al., 2021). These synthetic peptides provided excellent control on the sample homogeneity and were used to study ADP-ribose hydrolysis by different erasers. In another study, Schuller et al. used a synthetic ring-opened ADP-ribosyl group linked to a biotinylated peptide to generate a robust binding assay based on AlphaScreen technology (Schuller et al., 2017). This modified ADP-ribosyl group was shown to be non-hydrolysable, so that this assay technology could be successfully applied to readers and erasers. Even though these methods were shown to work robustly, the reagents used may not be easily accessible.

In proteins, many different residues can serve as acceptors of ADP-ribose. Residues such as serine, aspartate or glutamate form an O-glycosidic bond and lysine, arginine or asparagine form an N-glycosidic bond with ADP-ribose. Additionally, the linkage via an S-glycosidic bond can be formed with cysteine residues (Cohen and Chang, 2018). While many different ADP-ribosyl-hydrolases exist and can remove ADP-ribose from O-or N-glycosidic bonds, to date there is no human enzyme reported able to reverse the S-glycosidic linkage (Voorneveld et al., 2021). We reasoned that a naturally occurring S-glycosidically linked ADP-ribosyl group may serve as non-hydrolysable ADP-ribosyl binding probe and therefore could be used to measure the interaction to both hydrolysing- and non-hydrolysing ADP-ribosyl binders.

To generate S-glycosidically linked ADP-ribosyl groups in controlled manner, we used S1 subunit of pertussis toxin from the bacterium *Bordetella pertussis*, which is known to efficiently catalyse the transfer of a single ADP-ribosyl group to a specific C-terminal cysteine residue in the α_i_ subunits of heterotrimeric G proteins (Gα_i_) (Ashok et al., 2020; Katada, 2012). We recombinantly produced proteins with a C-terminal 10-mer peptide of Gα_i_ and still observed efficient modification. This allows, in theory, site-specific addition of ADP-ribose to any protein with accessible C-terminus. We used this system to generate a MARylated YFP protein with stable S-glycosidic bond able to bind ADP-ribosyl readers or erasers. We were able to further extend this system by using PARP enzymes that extend the single residue linked MAR to PAR, allowing us to probe the binding of both MAR and PAR binders. The *in vitro* system allows for simple and efficient setup of binding assays for ADP-ribosyl readers and erasers based on site-specific cysteine ADP-ribosylation of a Gα_i_-based tag we termed GAP (**G**i protein **A**lpha subunit **P**eptide). We further demonstrated the possibility to modify GAP-tagged proteins with chemically modified NAD^+^-analogs. To complement the binding assays, we developed a fast and simple detection method of mono- or poly-ADP-ribosylation on blots by fusing ADP-ribosyl binders to nanoluciferase (Nluc). These methods open ways for the development of various *in vitro* assay systems (**Figure 1**). To show the applicability for screening, we set up a binding assay for the macrodomain of SARS-CoV-2 non-structural protein 3 and identified the drug suramin as a moderate inhibitor.

**Figure 1:**
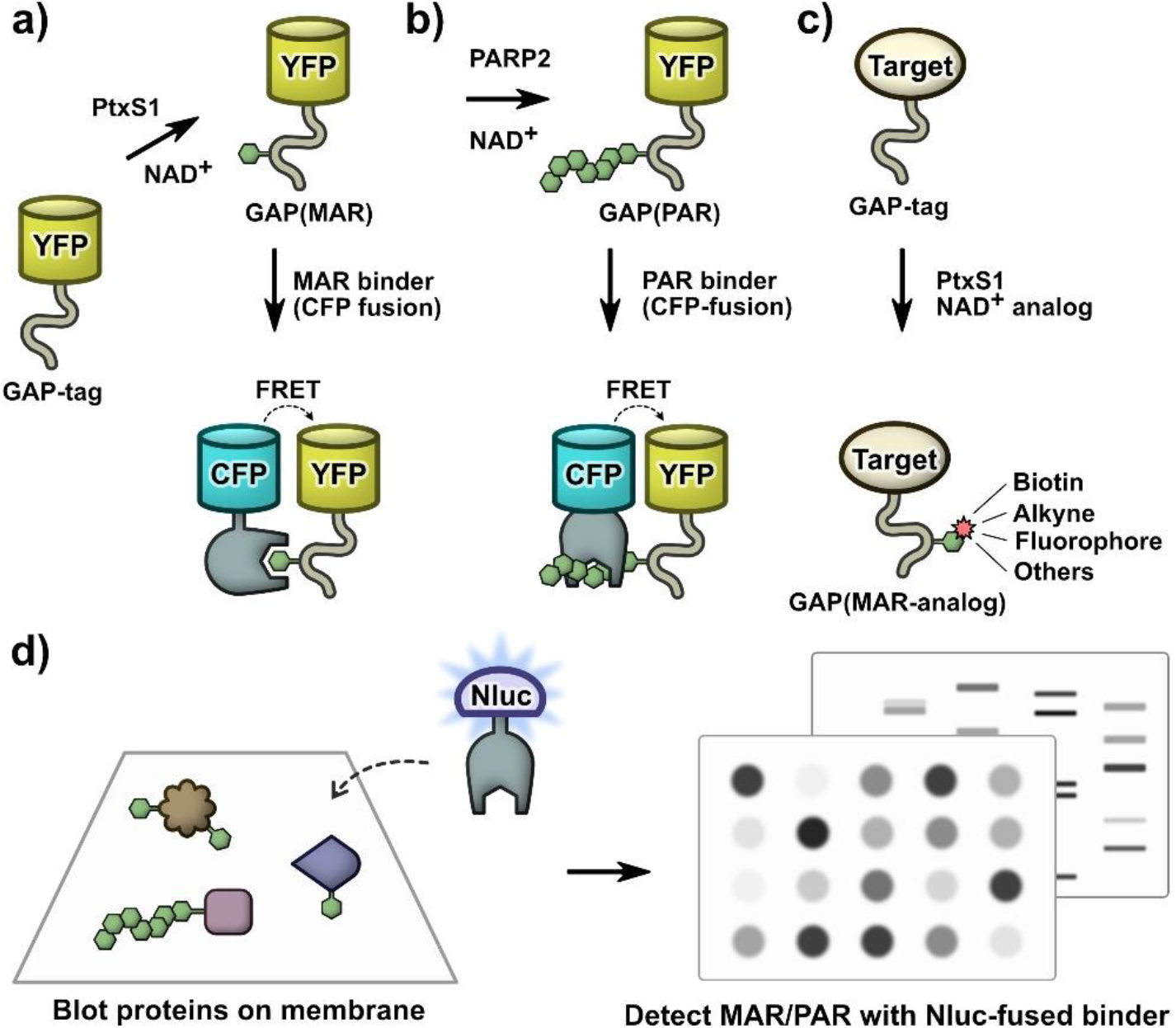
A molecular toolbox for *in vitro* interaction studies and assay development of ADP-ribosyl binding proteins. (a) Site-specific ADP-ribosylation of a C-terminal Gα_i_-based 10-mer peptide (GAP-tag) by pertussis toxin subunit S1 (PtxS1) allows for generation of single S-glycosidically linked mono-ADP-ribosyl (MAR) groups. (b) The MAR group of the GAP-tag can be extended to a poly-ADP-ribosyl (PAR) group by PARP2. This system can be used to measure binding of proteins interacting with mono- or poly-ADP-ribosyl groups by FRET or other binding technologies. (c) The GAP-tag can be used for site-specific labelling with NAD^+^ analogs. (d) High-affinity ADP-ribosyl binders fused to nanoluciferase (Nluc) can be used as luminescent probes for fast, sensitive, and selective detection of mono- and poly-ADP-ribosylated proteins in blot-based methods.

## Results

### Initial preparation of proteins for toolbox studies

The proteins used in this study were recombinantly produced in *E. coli*. We reasoned that mono-ADP-ribosylation (MARylation) of Gα_i_ by pertussis toxin would provide a good probe to test binding to ADP-ribosyl groups because the modification is well-defined at a single residue and is linked to cysteine via an S-glycosidic linkage. We initially assumed this linkage to be stable against enzymatic hydrolysis by many of the erasers, which was later during our work also experimentally confirmed by others (Voorneveld et al., 2021). It was previously shown that the recombinantly produced pertussis toxin subunit S1 (PtxS1) could be used to ADP-ribosylate Gα_i_ proteins *in vitro* (Ashok et al., 2020). We tested the ability of PtxS1 to modify unlabelled and YFP-fused full-length Gα_i_ as well as a 10-residue C-terminal peptide of Gα_i_ (GAP) when fused to YFP (**Figure 2a**). While ADP-ribosylation by PtxS1 was confirmed for these Gα_i_ constructs, the lower signal for the GAP-tag indicates less efficient modification by PtxS1 compared to full-length constructs. Similar was reported recently for synthetic Gα_i_ peptides (Eskonen et al., 2020) and further confirmed by NAD^+^-consumption assay (**Figure S1**). Despite this, we found that ADP-ribosylation activity by PtxS1 was sufficient to produce MARylated GAP-tagged YFP (**Figure S2**).

**Figure 2:**
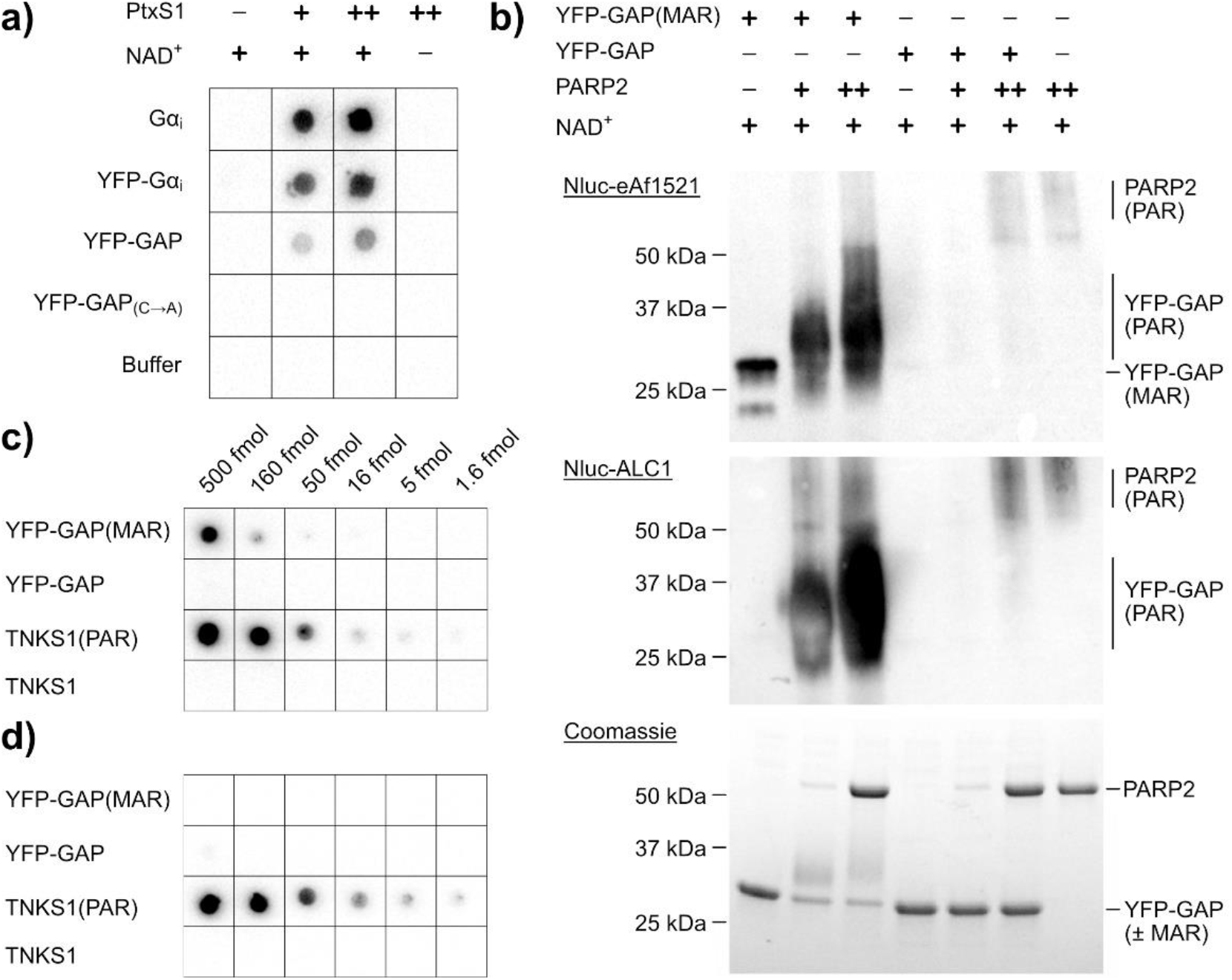
Initial development and preparation of toolbox components. (a) Testing ADP-ribosylation by PtxS1 with different Gα_i_ constructs. Unlabelled or YFP-fused full length Gα_i_ constructs and GAP-tagged YFP were tested as cysteine-ADPr acceptors when treated with 50 nM (+) or 250 nM (++) PtxS1. As controls, buffer or YFP-GAP in which the acceptor cysteine was mutated to alanine were used. Reactions were blotted on a nitrocellulose membrane and detection was done using Nluc-eAf1521. (b) The mono-ADP-ribosyl group in the GAP-tag can be extended to poly-ADP-ribose by PARP2. 10 µM YFP-GAP or YFP-GAP(MAR) were mixed with 1 mM NAD and 400 nM (+) or 4 µM (++) PARP2 or buffer as control. The reactions were run on SDS-PAGE and visualized using coomassie blue or by western-blot and detection using Nluc-eAf1521 or Nluc-ALC1. (c) Detection of MAR and PAR by Nluc-eAf1521. (d) Selective detection of PAR by Nluc-ALC1. Dilution series of YFP-GAP(±MAR) or TNKS1(±PAR) were blotted on nitrocellulose membranes. YFP-GAP: 1 fmol = 31 pg, TNKS1: 1 fmol = 75 pg.

Next, we aimed to introduce a poly-ADP-ribosyl (PAR) chain to the GAP-tag in order to generate a probe that could be used to detect interaction of PAR-binding proteins. We reasoned that the mono-ADP-ribosyl group of the previously modified GAP tag could serve as the starting point for elongation to poly-ADP-ribose by PARP enzymes. Detection of poly-ADP-ribosyl chains by western blot shows that the MARylated GAP-tag, but not the unmodified GAP-tag, can be PARylated by PARP2 in the presence of NAD^+^ (**Figure 2b, Figure S3**). A similar extension can be done by using TNKS1 instead of PARP2 (**Figure S4**).

As a tool to detect ADP-ribosylation, we adapted a method for blot-based detection by Nluc luminescence (Boute et al., 2016). Instead of fused antibodies, we produced the recently reported engineered ADP-ribosyl superbinder eAf1521 (Nowak et al., 2020) or high affinity PAR binder ALC1 (Singh et al., 2017) as fusion proteins with Nluc. We found that these constructs are easy to produce in *E. coli* and work in a simple and fast protocol for sensitive detection of mono- and poly-ADP-ribosylated proteins (**Figure 2c, d**).

### The GAP-tag for site-specific labelling of proteins using NAD^+^ analogs

While we could use the GAP-tag to site-specifically label proteins with ADP-ribose, we sought to demonstrate that this system could be extended with NAD^+^ analogs to introduce various chemical groups to the C-terminus of proteins. Many NAD^+^ analogs already exist and are commercially available such as biotinylated, fluorescent and click-chemistry ready NAD^+^ analogs (Depaix and Kowalska, 2019). We first tested the modification of GAP-tagged YFP with 6-biotin-17-NAD^+^ and confirmed that this NAD^+^ analog serves as substrate for PT-based modification of the GAP-tag (**Figure 3a**). The site-specific modification was detected using dot blot with streptavidin-conjugated horseradish peroxidase. We further tested modification of the GAP-tag with 6-propargyladenine-NAD^+^. The NAD^+^ analog is accepted as a substrate by PtxS1 and, as the resulting ADP-ribosyl-group contains an alkyne residue, it can be used in a copper(I)-catalyzed alkyne-azide cycloaddition reaction with azide-labelled Cy3 or Cy5 fluorophores to label the proteins (**Figure 3b**).

**Figure 3:**
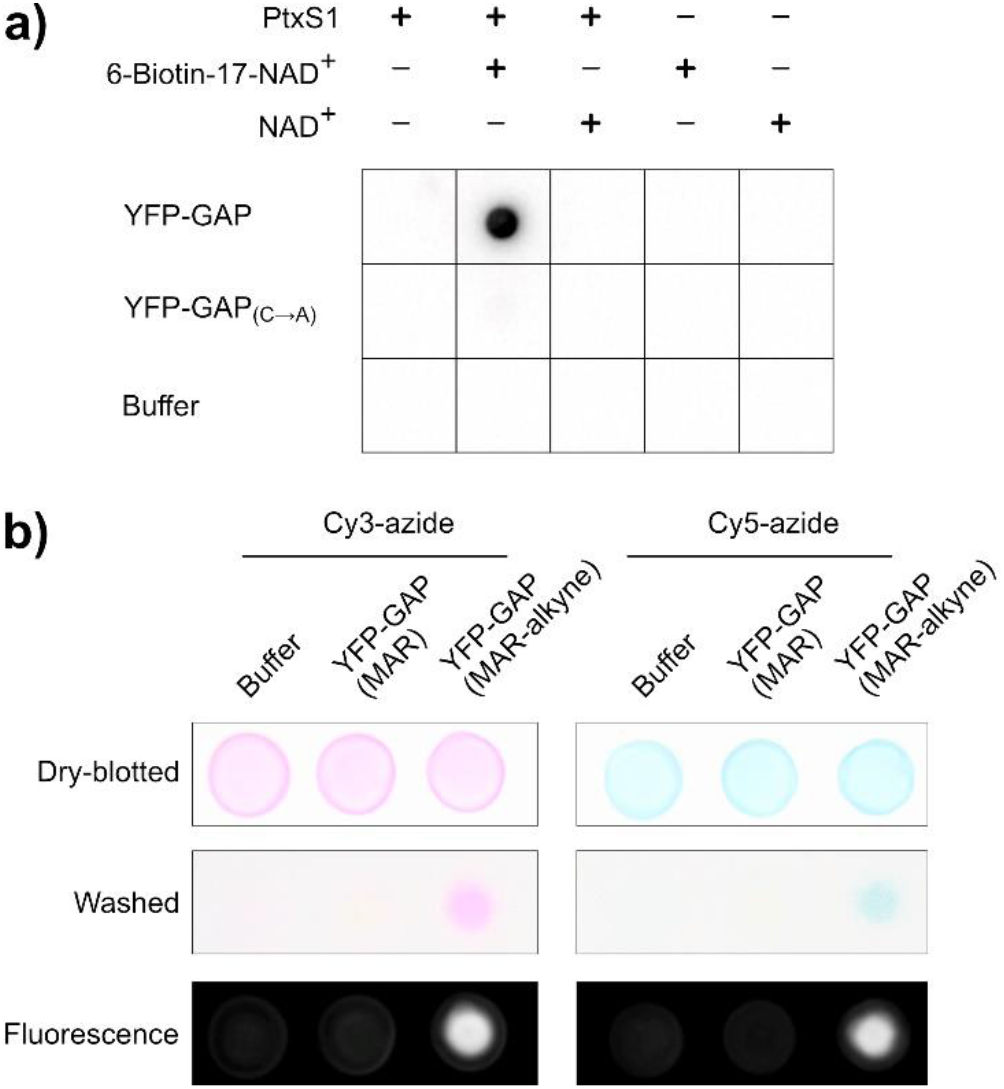
The GAP-tag can be used to introduce site-specific modifications with NAD^+^ analogs. (a) Site-specific biotinylation of the GAP-tag. GAP-tagged YFP was mixed with NAD^+^ or 6-Biotin-17-NAD^+^ in absence or presence of PtxS1. The reactions were blotted on a nitrocellulose membrane and detection of biotin was done with Streptavidin-HRP. (b) YFP-GAP was MARylated with PtxS1 using NAD^+^ or 6-propargyladenine-NAD^+^ containing an alkyne group. The resulting proteins YFP-GAP(MAR) or YFP-GAP(MAR-alkyne) or buffer were mixed with Cy3-azide or Cy5-azide and the copper(I)-catalyzed alkyne-azide cycloaddition reaction was performed by addition of 5 mM sodium ascorbate, 300 µM CuSO_4_ and 600 µM L-histidine. The samples were incubated for 3 hours at room temperature and blotted on nitrocellulose membranes and color images were taken. Unreacted Cy3-azide or Cy5-azide were removed by washing of the membranes in TBS-T. Color images as well as fluorescent images were taken.

### Evaluation of binding with confirmed and potential ADP-ribosyl binders

We selected a set of 27 different proteins to be tested for binding to the MARylated GAP-tag. We recombinantly produced these proteins in *E. coli* as fusions with CFP and tested ratiometric FRET signals upon binding to the MARylated YFP-GAP construct (**Figure 4a**). We used non-MARylated YFP-GAP construct as a control as well as MARylated YFP-GAP containing 200 µM ADP-ribose to compete with the interaction. A higher FRET signal correlates with a higher occupancy and thus binding-affinity of the binding partners, but it is also affected by the distance and orientation of the fluorophores (Kashida et al., 2017). We found that the proteins previously reported to bind ADP-ribose showed a higher FRET signal compared to the controls, indicating the binding to the MARylated GAP-tag. ARH1 was reported to bind ADP-ribose with a low affinity (Rack et al., 2018), which is likely the reason that we could not measure a FRET signal for this construct. While all three histone macrodomains macroH2A1.1, macroH2A1.2 and macroH2A2 have highly similar sequences, previous reports have concluded that only macroH2A1.1 has the ability to bind ADP-ribose (Kozlowski et al., 2018; Kustatscher et al., 2005). This is in agreement with our observations, wherein only macroH2A1.1 shows an elevated FRET signal over the controls. The GDAP2 protein has a macrodomain with similarities to MDO1 and MDO2, however it was reported to be unable to bind ADP-ribose (Neuvonen and Ahola, 2009), which is consistent with our findings.

**Figure 4:**
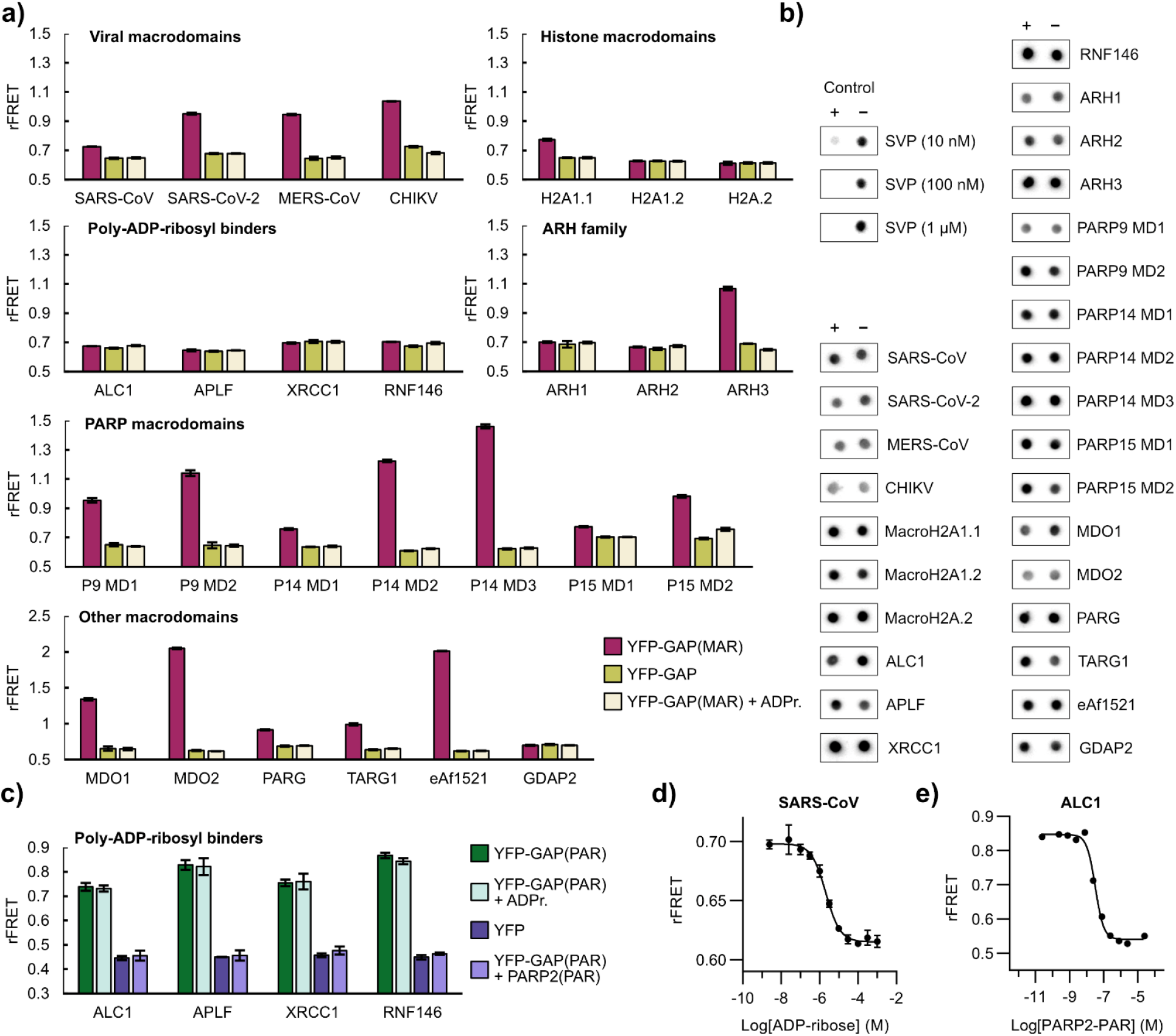
Testing interactions of reported and potential readers and erasers with YFP-GAP. (a) Interactions of CFP-fused potential and confirmed ADP-ribosyl binders with MARylated YFP-GAP. 1 µM CFP-fusion proteins were mixed with 5 µM YFP-GAP or with 5 µM YFP-GAP(MAR) in absence or presence of 200 µM ADP-ribose. The ratiometric FRET signals were measured. P9, P14, P15 = PARP9, PARP14, PARP15. (b) Test of ADP-ribosyl removal from GAP tag. 10 µM YFP-GAP(MAR) were prepared in absence (-) or presence (+) of 1 µM CFP-fused proteins or 0.01 µM to 1 µM snake venom phosphodiesterase I (SVP). Samples were incubated for 24 hours at room temperature and blotted on a nitrocellulose membrane. Detection was done with Nluc-eAf1521. (c) Interactions of poly-ADP-ribosyl binders with PARylated YFP-GAP. 250 nM CFP-fusion proteins were mixed with 500 nM YFP or with 500 nM YFP-GAP(PAR) in absence or presence of 100 µM ADP-ribose or 2.5 µM automodified PARP2. The ratiometric FRET signals were measured. (d) Representative dose-response curve of 1 µM CFP-SARS-CoV nsp3 and 5 µM YFP-GAP(MAR) upon competition with ADP-ribose. The control containing no ADP-ribose was set one logarithmic unit below the lowest concentration. (e) Representative dose-response curve of 250 nM CFP-ALC1 and 500 nM YFP-GAP(PAR) upon competition with PARylated PARP2. The control containing no PARP2(PAR) was set one logarithmic unit below the lowest concentration, while the control using YFP-GAP(MAR) instead of YFP-GAP(PAR) was set one logarithmic unit above the highest PARP2(PAR) concentration. Data shown are mean ± standard deviation with number of replicates n = 4.

While cysteine-(ADP-ribosyl)hydrolase activity was detected in human erythrocytes and mitochondria (Herrero-Yraola et al., 2001; Tanuma and Endo, 1990), no specific enzymes in humans with this activity could be identified to date. To test if any of the proteins show S-glycosylhydrolase activity, we mixed the CFP-fused constructs from above with MARylated YFP-GAP protein and tested hydrolysis of ADP-ribose using dot blot analysis after a 24-hour incubation period (**Figure 4b**). While the controls including snake venom phosphodiesterase I (SVP) showed loss of the signal through cleavage of the ADP-ribose diphosphate, none of the tested proteins were able to substantially hydrolyse the S-glycosidic bond under the conditions tested. ARH family members also showed no hydrolysis upon addition of MgCl2 (**Figure S5**). These findings highlight the versatility of this system for measuring the ADP-ribosyl binding of proteins that can otherwise hydrolyse O- or N-glycosidic linkages such as MDO1, MDO2, TARG1, ARH3 or SARS-CoV-2 nsp3.

The PAR binders ALC1, APLF, XRCC1 and RNF146 WWE domain did not show binding to MARylated YFP-GAP (**Figure 4a**), however they showed increased FRET signals when we used YFP-GAP PARylated by PARP2 (**Figure 4c**). For proteins binding to the MARylated GAP-tag, even comparatively low rFRET values showed good signals in dose-response experiments by competition with ADP-ribose (**Figure 4d, Figure S6**). Similarly, a representative curve for ALC1 with PARylated YFP-GAP was recorded and shows loss of the FRET signal with higher concentrations of auto-PARylated PARP2 protein (**Figure 4e**).

We used FRET for testing the interaction of proteins with the ADP-ribosylated GAP-tag, however we sought to demonstrate that binding to MARylated Gα_i_ could also be measured with different binding technologies. We used MDO2 as an example protein for this. Similarly to the binding of CFP-fused MDO2 to MARylated YFP-GAP to measure FRET (**Figure 5a, Figure S7**, Kd=654 nM), we used Nluc-fused MDO2 to generate a BRET signal upon interaction with MARylated YFP-GAP (**Figure 5b**). Conveniently, the same YFP variant can serve as acceptor in FRET and BRET applications. AlphaScreen protocols for ADP-ribosyl readers or erasers exist (Ekblad et al., 2018; Haikarainen et al., 2018; Schuller et al., 2017) and we could use our system adapted to AlphaScreen technology to directly probe binding to MARylated Gα_i_ by MDO2 (**Figure 5c**). We further showed binding of MDO2 to MARylated YFP-GAP using biolayer interferometry (BLI, **Figure 5d**), which in recent years has gained popularity as method for screening of small molecule compounds (Kaminski et al., 2017; Overacker et al., 2021; Peltomaa et al., 2018). This method also allows for simple quantification of kinetic binding parameters such as the dissociation constant (**Figure S8**).

**Figure 5:**
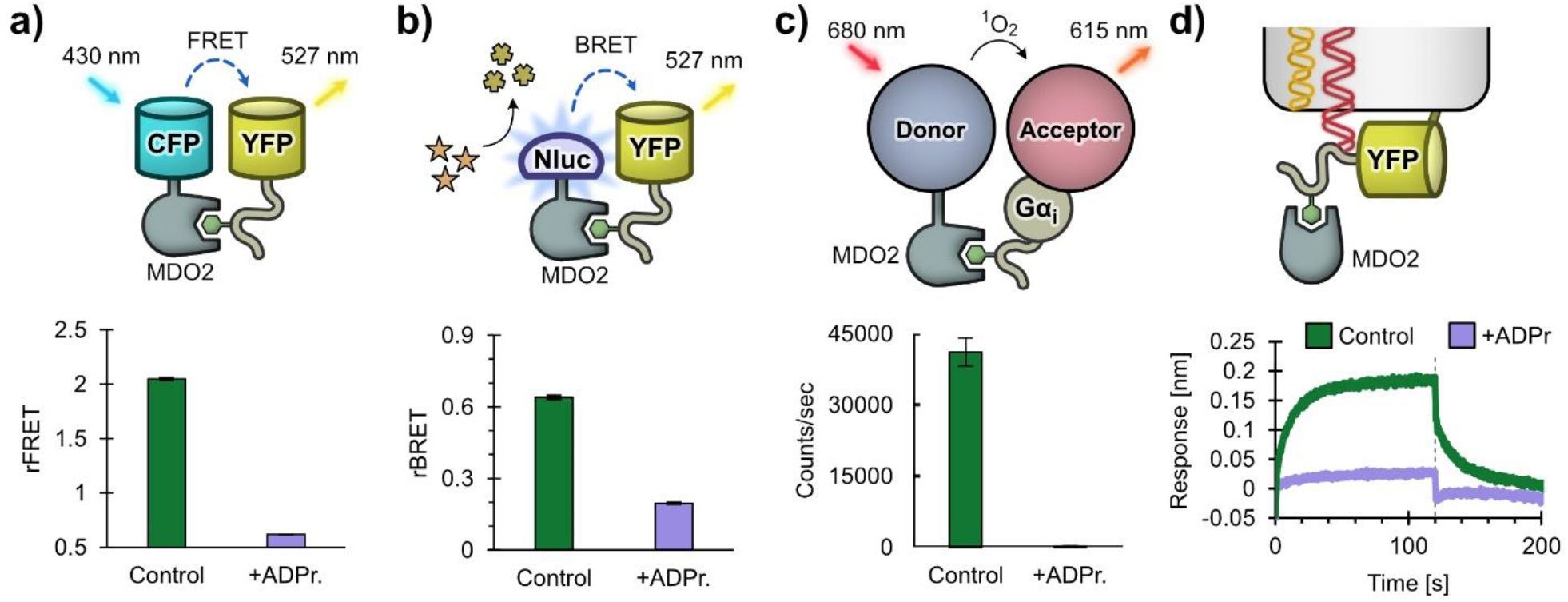
Various assay technologies can be utilized to detect binding to the MARylated Gα_i_. (a) Measurement of interaction by FRET. Ratiometric FRET signal of CFP-MDO2 and YFP-GAP(MAR) in absence (control) or presence of 200 µM ADP-ribose as shown in Figure 4a. (b) Measurement of interaction by BRET. Ratiometric BRET signal of Nluc-MDO2 and YFP-GAP(MAR) in absence (control) or presence of 200 µM ADP-ribose. (c) Measurement of interaction by AlphaScreen. Biotinylated MDO2 and His-tagged MARylated Gα_i_ were mixed with streptavidin donor beads and chelate acceptor beads in absence (control) or presence of 10 µM ADP-ribose. The luminescence signal was detected upon excitation of donor beads. (d) Measurement of interaction by biolayer interferometry. His-tagged YFP-GAP(MAR) was bound to the optical sensor surface and the change of signal after association (0 sec) or dissociation (120 sec, dotted line) of unlabelled MDO2 protein was determined in absence or presence of 3.16 µM ADP-ribose. Data shown in a-c are mean ± standard deviation with number of replicates n = 4.

### Application example: Screening for inhibitors against the SARS-CoV-2 nsp3 macrodomain

In light of the current situation regarding the COVID-19 pandemic, we chose to demonstrate the applicability of this binding system for screening of small molecule inhibitors against the macrodomain of SARS-CoV-2 nsp3. Presently, efforts are being made by researchers worldwide to find inhibitors against this macrodomain (Bonfiglio et al., 2020; Brosey et al., 2021; Cantini et al., 2020; Michalska et al., 2020; Ni et al., 2021; Russo et al., 2021; Schuller et al., 2021; Virdi et al., 2020). The nsp3 macrodomain of coronaviruses was shown to be critical for the viral replication (Fehr et al., 2015, 2016), and therefore small molecule inhibitors might show promise as therapeutic agents to fight infections caused by SARS-CoV-2 (COVID-19) and other viruses.

We assessed the quality of the FRET signals of the alternating positive and negative controls in a 384-well plate by mixing CFP-fused SARS-CoV-2 nsp3 macrodomain with MARylated YFP-GAP in the absence and presence of 200 µM ADP-ribose (**Figure 6a**). While the SARS-CoV-2 macrodomain was reported to have a relatively low binding affinity to ADP-ribose (Kd = 17 µM, Alhammad et al., 2021), we still found that the FRET signal showed sufficient separation of positive and negative controls. The Z’-factor was calculated to be 0.7, indicating that this assay is well suitable for high-throughput screening (Zhang et al., 1999).

**Figure 6:**
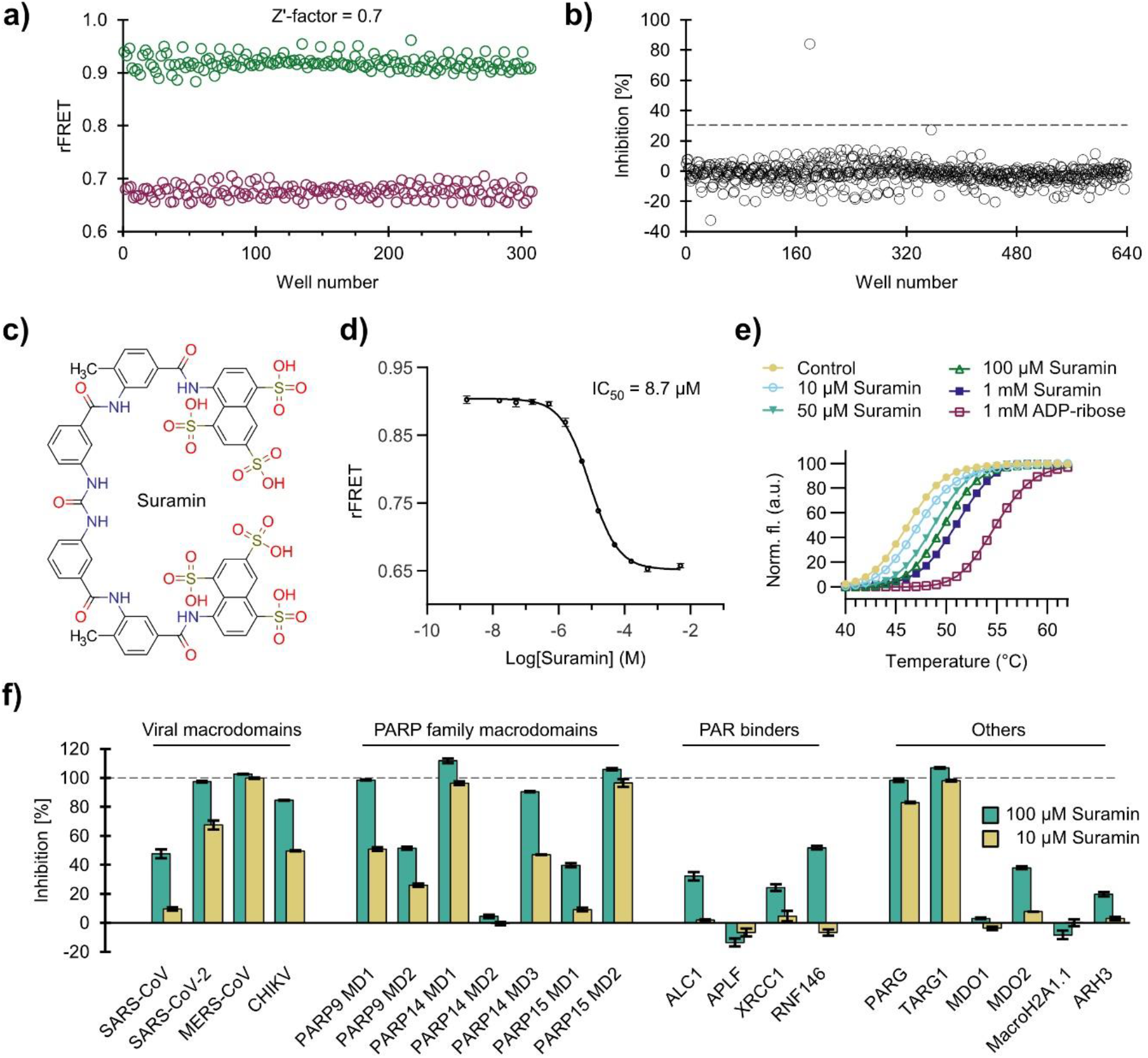
Development of a screening assay for the SARS-CoV-2 nsp3 macrodomain. (a) Signal validation for a screening assay with CFP-SARS-CoV-2. 1 µM SARS-CoV-2 was mixed with 5 µM YFP-GAP(MAR) in the absence (negative control) or presence (positive control) of 200 µM ADP-ribose and a Z’-factor of 0.7 was calculated. (b) Screen of ENZO FDA-approved drug library. One compound showed inhibition above 30% and was taken to further validation. (c) Structure of the hit compound suramin. (d) Dose-response curve with suramin shows an IC_50_ of 8.7 µM for the SARS-CoV-2 nsp3 macrodomain in the FRET-based. The control containing no compound was set one logarithmic unit below the lowest concentration, while the control containing 200 µM ADP-ribose was set one logarithmic unit above the highest suramin concentration. (e) Suramin shows stabilization of SARS-CoV-2 nsp3 macrodomain by DSF. (f) Inhibition profile of suramin against human and viral ADP-ribosyl binders used in this study. The inhibition was calculated based on the ratiometric FRET signals of the CFP-fused binders mixed with YFP-GAP(MAR) or YFP-GAP(PAR). Data shown are mean ± standard deviation with number of replicates n = 4.

We screened against an FDA-approved drug library comprising 640 small molecule compounds at 20 µM compound concentration (**Figure 6b**). From the screening, only the compound suramin was regarded as hit with 82% inhibition (**Figure 6c**). The IC50 value of this compound was determined to be 8.7 µM against the SARS-CoV-2 nsp3 macrodomain in the FRET-based assay (**Figure 6d**). To confirm binding of the compound to the SARS-CoV-2 nsp3 macrodomain, we performed nanoDSF analysis and showed stabilization of the protein by suramin in a concentration dependent manner (**Figure 6e**). Intriguingly, suramin is used as a broadband antiviral and antiparasitic drug and was recently reported to inhibit SARS-CoV-2 infection in cell culture-based models (Salgado-Benvindo et al., 2020) and to bind with high affinity to the RNA polymerase of SARS-CoV-2 (Yin et al., 2021). After reviewing the literature associated with suramin, we found that it is reported to inhibit a plethora of protein targets such as DNA- and RNA-polymerases, sirtuins, ATPases and G protein-coupled receptors (Freissmuth et al., 1996; Torrente et al., 2014; Trapp et al., 2007; Wiedemar et al., 2020), indicating that it exhibits low target specificity. We next tested the inhibition of suramin against the viral and human ADP-ribose readers and erasers produced in this study using the FRET-based assay (**Figure 6f**). Not surprisingly, suramin showed strong inhibition even at 10 µM against many of the proteins tested, confirming the low target specificity of this compound. To our knowledge, inhibition of macrodomains by suramin was not previously reported. We show that suramin also inhibits the nsp3 macrodomain of CHIKV, and inhibition of CHIKV pathogenesis by suramin was shown in multiple studies (Albulescu et al., 2015, 2020; Henß et al., 2016; Ho et al., 2015; Kuo et al., 2016; Lu et al., 2017). While multiple mechanisms for this inhibitory activity against CHIKV were reported, it is tempting to speculate that the inhibition of the nsp3 macrodomain poses another yet overlooked mechanism.

## Discussion

The high complexity of ADP-ribosyl associated pathways makes the involved proteins notoriously hard to study (Bonfiglio et al., 2020; Lüscher et al., 2018). While potent and specific inhibitors against many of the ADP-ribosyl transferases fundamentally helped to broaden our understanding of these enzymes (Durkacz et al., 1980; Huang et al., 2009; Kirby et al., 2018; Venkannagari et al., 2016; Wang et al., 2018), such inhibitors against ADP-ribosyl readers and erasers are still scarcely available or in early stages of development (Harrision et al., 2020; James et al., 2016; Liu et al., 2020; Palazzo and Ahel, 2018; Schuller et al., 2017). It was suggested that a possible bottleneck for the discovery of inhibitors is due to the lack of accessible high-throughput technologies (Schuller et al., 2017). Many of the assay systems work for only a subset of mono- or poly-ADP-ribosyl readers or erasers, require expensive or custom-made reagents or are not suited for high-throughput screening.

We have shown that the MARylated GAP-tag provides an easy to use and stable template for the development of binding assays for ADP-ribosyl binding proteins. This system can be adapted to different binding technologies such as FRET, BRET, AlphaScreen or BLI as shown in this study – but is not limited to these and could be extended to work with technologies such as protein-fragment complementation assays or time resolved-FRET methods. The GAP-tag can also be used to site-specifically introduce chemical ADP-ribose analogs to the protein of interest, adding another technology to the chemical biology toolkit of protein labelling tags (Lotze et al., 2016). The ADP-ribosyl labelling technology ELTA might further be used to extend this functionality (Ando et al., 2019). We have further shown that the mono-ADP-ribosyl group attached to the GAP-tag can be subsequently extended to poly-ADP-ribosyl chains by PARP2, which we used to detect interactions with PAR binding proteins. A similar strategy for generation of defined single poly-ADP-ribosyl chains to protein targets was to our knowledge not reported before and might prove useful also in other applications in the future.

We tested in this study a total of 27 proteins that were either confirmed to bind to ADP-ribosyl groups or were shown not to bind them, despite being macrodomains. For 22 of these proteins, we confirmed binding to either mono- or poly-ADP-ribosyl groups attached to the GAP tag. The remaining 5 proteins not showing binding were either reported not to bind ADP-ribose or bind it only with very low affinity. The systems described in this work can be entirely recombinantly produced in *E. coli* with good yields for most of the proteins (>200 mg/L culture). While we have produced and tested a large set of proteins in this study, the production of only three proteins is required to set up a high-throughput assay for an ADP-ribosyl binder of interest (for instance: CFP-fusion protein, YFP-GAP and PtxS1). In the frame of our work, we also made use of Nluc-fused high-affinity ADP-ribose binders eAf1521 and ALC1 as sensitive detection agents of mono-and poly-ADP-ribosyl groups for blot-based studies, respectively. Gibson et al. described a similar system based on ADP-ribosyl binding proteins fused to the Fc region of rabbit immunoglobulin, which can subsequently be detected by commercial antibodies targeting the Fc region (Gibson et al., 2017). Due to simple recombinant expression, we used Nluc and not the more established horse radish peroxidase as luminescent reporter. This may require more optimization efforts for quantitative blots, but it however works readily for semi-quantitative or qualitative detection. We found that our system, which does not require a secondary antibody, takes only little working time of about 1 hour from blotting to imaging in practice. We used these to confirm mono- or poly-ADP-ribosylation of the GAP tag and other Gα_i_ constructs and to test for potential hydrolysis of S-glycosidic bonds by the proteins used in this study, but it could also be used for studying ADP-ribosylation by PARP family members or might find applications in cell-based studies.

As an example for an application of this GAP-based binding assay, we screened a small molecule library of existing drugs against the nsp3 macrodomain of SARS-CoV-2. We validated suramin as a hit compound, confirming that the assay system is suitable for screening of inhibitors. While suramin showed significant inhibition of other ADP-ribosyl binders utilizing the same assay system, we believe that the simple setup of profiling of potential inhibitors against many different ADP-ribosyl binders in itself is a valuable application of this technology.

In summary, the tools presented in this study are very accessible and allow setting up of robust and adaptable ADP-ribosyl binding assays and will therefore aid in the investigation of ADP-ribosyl binders and the discovery of chemical probes targeting them.

## Methods

### Cloning

Detailed cloning procedures are described in the supplementary material. Boundaries of the cloned constructs are shown in **Supplementary Table S1**. Briefly, expression constructs were cloned into expression vectors based on pNIC28-Bsa4 or pNH-TrxT by SLIC (Jeong et al., 2012).

### Protein expression

Detailed protein expression procedures are described in the supplementary material. Briefly, *E. coli* BL21(DE3) or *E. coli* Rosetta 2 cells were transformed with the plasmids encoding the expression constructs. Terrific Broth (TB) autoinduction media including trace elements (Formedium, Hunstanton, Norfolk, England) was supplemented with 8 g/l glycerol and antibiotics and inoculated with 1:100 of preculture grown over night in LB. The flasks were incubated shaking at 37 °C until an OD600 of about 1 was reached. The temperature was set to 15-18 °C and incubation continued overnight. The cells were collected by centrifugation at 4,200×g for 15-30 min at 4 °C. The pellets were thereafter resuspended in lysis buffer. Resuspended cells were stored at −20 °C until purification.

### Protein purification

Detailed protein expression procedures are described in the supplementary material and **Supplementary Table S2**. Briefly, the cells were thawed and lysed by sonication and all constructs were initially purified by immobilized metal affinity chromatography (IMAC). Following IMAC, an additional size exclusion purification was performed for most of the proteins. Finally, proteins were concentrated and subsequently aliquoted and flash frozen in liquid nitrogen and stored at −70 °C.

### Preparation of mono- and poly-ADP-ribosylated GAP-tagged proteins

YFP with C-terminal GAP-tag was purified by IMAC and dialyzed against 20 mM HEPES pH 7.5, 350 mM NaCl. YFP-GAP was diluted to 100 µM in 50 mM sodium phosphate buffer pH 7.0 and mixed with 1.5 µM PtxS1 and 150 µM NAD^+^. The reaction was incubated for an hour at room temperature. To ensure completeness of the reaction, a second 150 µM were added to the reaction. Incubation was continued for 1 h at room temperature. The reaction mixture loaded to an IMAC column to remove pertussis toxin, hydrolysis products and unreacted NAD^+^. IMAC was carried out as described in the purification procedures above. The buffer was exchanged to 20 mM HEPES pH 7.5, 150 mM NaCl, 0.5 mM TCEP and the MARylated YFP-GAP was subsequently concentrated to about 1 mM concentration using a Amicon Ultra-15 Centrifugal Filter Unit (MWCO: 10kDa). The protein was flash frozen in liquid nitrogen and stored at −70 °C.

PARylated YFP-GAP for FRET experiments was prepared from MARylated YFP-GAP. 10 µM of MARylated YFP-GAP was incubated in the presence of 400 nM PARP2 (residues 90-583) and 1 mM NAD^+^ in a buffer solution containing 50 mM Tris pH 8.0 and 5 mM MgCl2 for 2 h at room temperature. The reacted sample was then purified using IMAC as described above to remove PARP2. The sample buffer was exchanged to 30 mM HEPES pH 7.5, 150 mM NaCl, 10% glycerol, 0.5 mM TCEP using an Amicon Ultra-15 Centrifugal Filter Unit (MWCO: 10kDa). The protein was aliquoted and flash frozen in liquid nitrogen and stored at −70 °C.

For Tankyrase1 PARylated YFP-GAP production, MARylated YFP-GAP was incubated with 200 nM Tankyrase 1 SAM-catalytic domain dimer and 1 or 10 mM NAD^+^ in a buffer solution containing 10 mM BisTrisPropane [pH7.0], 0.01% Triton X100. The reaction has carried out for 16 h at room temperature.

### Blot-based detection of mono- and poly-ADP-ribosylation

For dot blot experiments, we transferred 0.5 µl per spot of the sample solution to dry nitrocellulose membranes using Echo 650. All following steps were performed at room temperature. After drying of the spots, the membrane was blocked on a shaker for 10 min in 15 ml 5%(w/v) skimmed milk powder in TBS-T. The blocking solution was discarded, and the membrane was incubated on a shaker with 15 ml of 0.1 µg/ml Nluc-eAf1521 or Nluc-ALC1 in 1%(w/v) skimmed milk powder in TBS-T. After discarding the Nanoluc-eAf1521 solution, the membrane was rinsed with 15 ml TBS-T and incubated on a shaker with 15 ml TBS-T for 15 min. After a final rinsing with 15 ml TBS-T, the membrane was imaged using 500 µl of 1:1000 NanoGlo substrate (Promega, catalogue number: N1120) diluted in 10 mM sodium phosphate buffer pH 7.0.

For western blot, 10 µl samples were first run in SDS-PAGE (Mini-Protean TGX 4-20% gradient gel, BioRad). The proteins were then transferred to a nitrocellulose membrane (Trans-Blot Turbo, BioRad) using TransBlot semi dry system (BioRad). After transfer, membranes were treated following the same procedure as described above for dot-blot. Nanoluc-eAF1521 and Nanoluc-ALC1 were used at 0.1 µg/ml.

### Testing Nluc-eAf1521 and Nluc-ALC1 sensitivity and selectivity

Dilution series of YFP-GAP, MARylated YFP-GAP, MBP-TNKS1 construct or auto-PARylated MBP-TNKS1 construct were blotted on a nitrocellulose membrane and detected with Nluc-eAf1521 or Nluc-ALC1 as described above.

To generate auto-PARylated TNKS1, 10 µM of purified TNKS1 construct were mixed with 1 mM NAD^+^ in 50 mM Bis-Tris-Propane pH 7.0, 0.01% Triton X-100, 0.5 mM TCEP. To generate a control containing TNKS1 without PAR, partial auto-PARylation that occurred during recombinant expression in *E. coli* was removed by mixing TNKS1 construct with 2 µM snake venom phosphodiesterase I from *Crotalus adamanteus* (Worthington Biochemical Corporation).

### Modification test of YFP-GAP with 6-Biotin-17-NAD^+^

10 µM YFP-GAP or YFP-GAP(cysteine to alanine mutant) were mixed with 1 µM NAD^+^ or 6-Biotin-17-NAD^+^ (Biolog). Reactions were prepared in absence or presence of 0.5 µM PtxS1 and were incubated for 1 h at room temperature and then blotted to a dry nitrocellulose membrane (0.5 µl per spot of the sample solution to dry nitrocellulose membranes using Echo 650). The membrane was let dry and thereafter blocked on a shaker for 10 min in 15 ml blocking buffer (1% casein in TBS, BioRad). The blocking solution was discarded, and the membrane was incubated on a shaker for 1 hour with 15 ml of 1:5000 Streptavidin-HRP in blocking buffer. After discarding the Streptavidin-HRP solution, the membrane was rinsed with 15 ml TBS-T and incubated on a shaker with 15 ml TBS-T for 15 min. After a final rinsing with 15 ml TBS-T, the membrane was imaged using ECL solution (BioRad).

### Modification test of YFP-GAP with 6-propargyladenine-NAD^+^ and addition Cy3 and Cy5 azides by CuAAC

YFP-GAP(6-propargyladenine-MAR) was prepared as described above for YFP-GAP using 6-propargyladenine-NAD^+^ instead of NAD^+^. To test the addition of Cy3 or Cy5 to YFP-GAP(6-propargyladenine-MAR) by CuAAC, reactions were prepared in 25 mM HEPES pH 7.5 by mixing 15 µM of YFP-GAP(6-propargyladenine-MAR) or YFP-GAP(MAR) with 5 mM sodium ascorbate, 50 µM Cy3-azide or Cy5-azide and pre-mixed 300 µM CuSO4 and 600 µM L-histidine. Additionally, controls without protein were prepared. The reactions were let incubate for 3 hours at room temperature and afterwards blotted on a nitrocellulose membrane (5 µl per spot). The membrane was washed in 15 ml TBS-T for 30 min and imaged. Fluorescence imaging was done with an Azure 600 imaging system (Azure Biosystems) using Cy3 or Cy5 filter settings, respectively.

### FRET measurement

The measurements were done as previously described (Sowa et al., 2020). Briefly, the samples were excited at 410 nm and emission at 477 nm and 527 nm wavelengths were measured. The ratiometric FRET value (rFRET) was calculated by dividing the fluorescence intensity at 527 nm by the fluorescence intensity at 477 nm. The experiments with MARylated YFP-GAP were carried out in assay buffer (10 mM Bis–Tris-Propane pH 7.0, 3% (w/v) PEG20,000, 0.01%(v/v) Triton X-100 and 0.5 mM TCEP) in 10 µl volume per well whereas those with PARylated YFP-GAP were done in 10 mM Tris pH 8.0, 150 mM NaCl, 0.01% Tween-20 in 20 µl, unless stated otherwise.

### BRET measurement

The reactions were performed in 384-well white OptiPlates (PerkinElmer). A reaction volume of 40 µl per well was used. 50 nM Nluc-MDO2 were mixed with 1 µM MARylated YFP-GAP. The reaction was started by addition of 1:4000 NanoGlo substrate (Promega, catalogue number N1110). The reaction was incubated for 5 minutes and the emission was measured at wavelengths of 445-470 nm and 520-545 nm using Tecan Spark multimode plate reader with luminescence readout and a settle time of 10 ms and integration time of 500 ms. The ratiometric BRET value (rBRET) was calculated by dividing the luminescence intensity and 520-545 nm by the luminescence intensity at 445-470 nm. The experiments were carried out in assay buffer (10 mM Bis–Tris-Propane pH 7.0, 3% (w/v) PEG20,000, 0.01%(v/v) Triton X-100 and 0.5 mM TCEP).

### AlphaScreen

AlphaScreen technology was utilized to demonstrate the assay principle as described previously (Haikarainen et al., 2018). The reaction was performed in a 384-well flat-grey Alphaplate (PerkinElmer) in a total volume of 25 µl. The reaction consisted of 300 nM His-tagged Gα_i_(MAR) mixed with 300 nM biotinylated MDO2 in a buffer containing 25 mM HEPES pH 7.5, 100 mM NaCl and 0.1 mg/ml BSA. The plate was sealed and incubated for 80 min at room temperature with constant shaking at 300 rpm. Finally, 5 µg/ml nickel chelate acceptor and streptavidin donor beads were added to the plates followed by additional 3 hrs incubation. The plate contained blank wells (assay buffer and AlphaScreen beads only), control 1 (biotinylated MDO2, His-tagged Gα_i_) and control 2 (biotinylated MDO2, MARylated His-tagged Giα and 10 µM ADP-ribose). Luminescence was read using Tecan infinite M1000 Pro plate reader with AlphaScreen detection module.

### Biolayer interferometry

Biolayer interferometry (BLI) assays were carried out in Octet Red system (ForteBio) in a buffer containing 10 mM Bis-Tris-Propane pH 7.0, 150 mM NaCl, 1% BSA, 0.02% Triton X-100 and at 30 °C and shaking at 1500 rpm. 10 µg/ml YFP-GAP or MARylated YFP-Gi was loaded on Ni^2+^-NTA coated sensors, followed by a wash step in buffer. Association to MDO2 was measured by dipping the sensors in solution containing 0-2 µM MDO2 for 120 s, while for the dissociation step the sensors were dipped in buffer for 120 s.

For the ADP-ribose competition experiments, 10 µg/ml YFP-GAP or MARylated YFP-GAP were loaded onto Ni^2+^-NTA coated sensors. For association, sensors were dipped in 100 nM MDO2 mixed with a half-log dilution series of ADPr (10 nM to 10 µM) for 120 s and then transferred to buffer for the dissociation step.

### Cysteine-ADP-ribosyl hydrolysis assay

All samples were incubated for 24 hours at room temperature prior to blotting. 1 µM of CFP-fused proteins or 0.01, 0.1 and 1 µM of SVP were mixed with 10 µM MARylated YFP-GAP in 10 mM HEPES pH 7.5, 25 mM NaCl, 0.5 mM TCEP. After incubation, 0.5 µl per spot of the reaction mixtures were blotted next to 0.5 µl spots of 10 µM MARylated YFP-GAP. As control, 10 µM of non-MARylated YFP-GAP was blotted next to 10 µM of MARylated YFP-GAP.

### Validatory screening

For the screening, 40 nl of 10 mM compound stocks dissolved in DMSO from the FDA-approved drug library (Enzo Life Sciences) were transferred to 384-well black low-volume polypropylene plates (Fisherbrand). The sample mixture containing 1 µM CFP-SARS-CoV-2 nsp3 macrodomain and 5 µM MARylated YFP-GAP was prepared in assay buffer (10 mM Bis–Tris-Propane pH 7.0, 3%(w/v) PEG20,000, 0.01%(v/v) Triton X-100 and 0.5 mM TCEP) and 20 µl per well were dispensed using Mantis liquid dispenser (Formulatrix). The rFRET signal was determined after 5-minute incubation time. The sample mixtures in presence or absence of 200 µM ADP-ribose were used as positive and negative controls, respectively.

### Differential scanning fluorimetry

The SARS-CoV-2 nsp3 macrodomain without tags was diluted to 5 µM in 10 mM HEPES pH 7.5, 25 mM NaCl, 0.5 mM TCEP buffer and mixed with 5x SYPRO Orange. Samples were prepared with 10 µM, 50 µM, 100 µM or 1 mM of suramin. Samples in presence or absence of 1 mM ADP-ribose were used as controls. Samples were transferred to 96-well qPCR plates. Measurement was performed in a BioRad C1000 CFX96 thermal cycler. Data points for melting curves were recorded in 1 min intervals from 20– 95 °C, with the temperature increasing by 1 °C/min. The analysis of the data was done in GraphPad Prism 7 using a nonlinear regression analysis (Boltzmann sigmoid equation) of normalized data.

## Data availability

The data that support the findings of this study are available from the corresponding author upon reasonable request. Expression constructs generated in this study will be available through Addgene.

## Acknowledgements

This work was funded by the Jane and Aatos Erkko and Sigrid Jusélius Foundations. The use of the facilities of the Biocentre Oulu Structural Biology core facility, member of Biocentre Finland, Instruct-ERIC Centre Finland and FINStruct, as well as of Proteomics and Protein Analysis and Sequencing core facilities are gratefully acknowledged. We thank Dr. Yashwanth Ashok for cloning of the pNH-Nluc vector and Jere Hukkanen for assisting with ARH1-3 protein production.

## Author information

Contributions

LL conceived the research. STS, AGP, SW, HIA and MMM cloned and produced the proteins used in the studies. STS and AGP developed the assay principles for MAR and PAR binders, respectively. STS developed the dot blot detection method and carried out inhibitor screening. AGP carried out the BLI assay and SW the AlphaScreen assay. SW did Kd measurements with the FRET assay. STS, AGP and LL wrote the publication with contributions from all the authors.

## Ethics declarations

### Competing interest

The authors declare the following competing financial interests: STS, AGP and LL are inventors listed in a patent application related to the described methods and these authors declare no additional
interests. The remaining authors declare no competing interests.

## Notes

### Summary of Updates

Author order corrected. Conflict of interest statement updated. Expression constructs will be deposited to Addgene and available through them.

