## Supplementary information for "A molecular toolbox for ADP-ribosyl binding proteins"

[#]These authors contributed equally

#### Content

**Expression vectors.**

**YFP-GAP sequence.**

**Table S1: Expression construct information.**

**Protein production procedures.**

**Table S2: IMAC purification information.**

**Figure S1: $NAD^+$ consumption by PtxS1 with different $G\alpha_i$ constructs.**

**Figure S2: Analysis of YFP-GAP ADP-ribosylation by mass spectrometry.**

**Figure S3: PARP2 can extend modify YFP-GAP-MARylated to produce a PARylated form.**

**Figure S4: PARylation of YFP-GAP by TNKS1.**

**Figure S5: Hydrolysis test of cysteine-ADP-ribose with ARH family members.**

**Figure S6: ADP-ribose dose-response curves for CFP-fusion constructs and MARylated YFP-GAP.**

**Figure S7: Determination of binding affinity for CFP-MDO2 and YFP-GAP(MAR) by FRET.**

**Figure S8: Use of BLI to measure MDO2 to YFP-GAP-MARylated interaction.**

**References**

#### Expression vectors

The expression vectors are based on pNIC28-Bsa4 (Addgene Plasmid #26103) or pNH-TrxT (Addgene Plasmid #26106). For pNIC-MBP, we inserted the sequence for *E. coli* maltose binding protein (UniProt ID: P0AEX9, residues 27-392) between $His_6$-tag and TEV protease cleavage site of pNIC28-Bsa4 as previously described (Sowa et al., 2020). In the same manner for pNIC-YFP and pNIC-CFP, the sequences encoding mCitrine (Addgene Plasmid #29724) or mCerulean (Addgene Plasmid #29726) were inserted between $His_6$-tag and TEV protease cleavage site of pNIC28-Bsa4. For pNH-Nluc, the sequence encoding Nluc (Addgene Plasmid #87696) was inserted into pNH-TrxT in place of the sequence encoding TrxT.

#### YFP-GAP sequence

Sequence for YFP-GAP cloned into pNIC28-Bsa4. The sequence of the 10-mer C-terminal Gai peptide is bold and underlined:

VSKGEELFTGVVPILVELDGDVNGHKFSVSGEGEGDATYGKLTLKFICTTGKLPVPWPTLVTTFGYGL
MCFARYPDHMKQHDFFKSAMPEGYVQERTIFFKDDGNYKTRAEVKFEGDTLVNRIELKGIDFKEDGNI
LGHKLEYNYNSHNVYIMADKQKNGIKVNFKIRHNIEDGSVQLADHYQQNTPIGDGPVLLPDNHYLSYQ
SKLSKDPNEKRDHMVLLEFVTAAGITLGMDELY**KNNLKDCGLF**

**Table S1: Expression construct information.**

| Construct name | Uniprot identifier | Boundaries (cloned into respective vector after ENLYFQSM) | Vector | Reference for cloning of the construct |
|---|---|---|---|---|
| CFP-ALC1 | Q86WJ1 | L613-K879 | pNIC-CFP | this work |
| CFP-APLF | Q8IW19 | K360-V448 | pNIC-CFP | this work |
| CFP-ARH1 | P54922 | E2-L357 | pNIC-CFP | this work |
| CFP-ARH2 | Q8NDY3 | E2-K354 | pNIC-CFP | this work |
| CFP-ARH3 | Q9NX46 | A14-S363 | pNIC-CFP | this work |
| CFP-CHIKV (nsp3) | A0A4D6GPC4 | A1334-T1493 | pNIC-CFP | this work |
| CFP-eAf1521 | O28751 | KH-M1-L192* | pNIC-CFP | this work |
| CFP-GDAP2 | Q9NXN4 | D1-N231 | pNIC-CFP | this work |
| CFP-MacroH2A1.1 | O75367-2 (isoform 1) | Q162-N369 | pNIC-CFP | this work |
| CFP-MacroH2A1.2 | O75367-1 (isoform 2) | Q162-N372 | pNIC-CFP | this work |
| CFP-MacroH2A.2 | Q9P0M6 | K167-K372 | pNIC-CFP | this work |
| CFP-MDO1 | Q9BQ69 | S58-A325 | pNIC-CFP | this work |
| CFP-MDO2 | A1Z1Q3 | K7-D243 | pNIC-CFP | this work |
| CFP-MERS-CoV (nsp3) | K9N7C7 | D1109-D1275 | pNIC-CFP | this work |
| CFP-PARG | Q86W56 | S448-T976 | pNIC-CFP | this work |
| CFP-PARP14 (MD1) | Q460N5 | K791-F978 | pNIC-CFP | this work |
| CFP-PARP14 (MD2) | Q460N5 | W1003-A1190 | pNIC-CFP | this work |
| CFP-PARP14 (MD3) | Q460N5 | F1208-G1388 | pNIC-CFP | this work |
| CFP-PARP15 (MD1) | Q460N3 | N78-S267 | pNIC-CFP | this work |
| CFP-PARP15 (MD2) | Q460N3 | G287-N470 | pNIC-CFP | this work |
| CFP-PARP9 (MD1) | Q8IXQ6 | G102-K298 | pNIC-CFP | this work |
| CFP-PARP9 (MD2) | Q8IXQ6 | N310-N493 | pNIC-CFP | this work |
| CFP-RNF146 (WWE) | Q9NTX7 | R99-L183 | pNIC-CFP | this work |
| CFP-SARS-CoV (nsp3) | P0C6X7 | E1000-K1173 | pNIC-CFP | this work |
| CFP-SARS-CoV-2 (nsp3) | P0DTD1 | E1084-E1192 | pNIC-CFP | this work |
| CFP-TARG1 | Q8IXB3 | A2-L152 | pNIC-CFP | this work |
| CFP-XRCC1 | P18887 | E315-P406 | pNIC-CFP | this work |
| Gαi (full length) | P63096 | G2-F354 | pNIC28-Bsa4 | Ashok et al., 2020 |
| MDO2 | A1Z1Q3 | K7-D243 | pNH-TrxT | Wazir et al., 2021 |
| NanoLuc-ALC1 | Q86WJ1 | L613-K879 | pNIC-Nluc | this work |
| NanoLuc-eAf1521 | O28751 | KH-M1-L192* | pNIC-Nluc | this work |
| NanoLuc-MDO2 | A1Z1Q3 | K7-D243 | pNIC-Nluc | this work |
| PARP2 (full-length) | Q9UGN5 | 2A-W583 | pNH-TrxT | Obaji et al., 2020 |
| PARP2 (WGR90-Cat) | Q9UGN5 | G90-W583 | pNH-TrxT | Obaji et al., 2020 |
| PtxS1 | P04977 | D35-I221 | pNIC28-Bsa4 | Ashok et al., 2020 |
| SARS-CoV-2 (nsp3) | P0DTD1 | E1084-E1192 | pNH-TrxT | this work |
| TNKS1 | O95271 | T1017-T1327 | pNIC-MBP | this work |
| TNKS1 (E1050K) | O95271 | T1017-Q1325, E1050K | pNIC-BSA4 | this work |
| TNKS1 (Y1073A) | O95271 | T1017-Q1325, Y1073A | pNIC-BSA4 | this work |
| YFP | P42212 | YFP: V1-Y237** | pNIC28-Bsa4 | Sowa et al., 2020 |
| YFP-GAP | YFP: P42212<br>Gαi: P63096 | YFP: V1-Y237**<br>Gαi: K345-F354 | pNIC28-Bsa4 | this work |
| YFP-GAP(C→A) | YFP: P42212<br>Gαi: P63096 | YFP: V1-Y237**<br>Gαi: K345-F354, C351A | pNIC28-Bsa4 | this work |
| YFP-Gαi (full length) | P63096 | M1-F354 | pNIC-YFP | this work |

* Point mutations correspond to the engineered Af1521 variant described by Nowak et al., 2020.

** Point mutations correspond to the YFP variant mCitrine found in Addgene plasmid #29771.

### Protein production procedures

Expression and purification procedures for full length G$\alpha_i$ (Ashok et al., 2020), MDO2 (Wazir et al., 2021), PARP2 constructs (Obaji et al., 2020) and YFP (Sowa et al., 2020) were performed as previously described.

#### Protein expression

The following constructs were expressed in *E. coli* strain Rosetta2(DE3): CFP-SARS-CoV, CFP-MacroH2A2, CFP-PARP9 (MD2), CFP-PARP14 (MD1), CFP-PARP14 (MD2), CFP-PARP14 (MD3), CFP-PARP15 (MD2), CFP-MDO2, CFP-PARG and CFP-GDAP2. All other constructs were expressed in *E. coli* strain BL21(DE3). The respective chemically competent cells were transformed with the plasmids described in **Table S1**. 500 ml Terrific Broth (TB) autoinduction media including trace elements (Formedium, Hunstanton, Norfolk, England) were supplemented with 8 g/l glycerol and 50 µg/ml kanamycin and inoculated with 5 ml of overnight preculture. The media was additionally supplemented with 34 µg/ml chloramphenicol for Rosetta2(DE3) cells. The flasks were incubated shaking at 37 °C until an $OD_{600}$ of about 1 was reached. The temperature was thereafter set to 15 °C for constructs PtxS1, CFP-PARP14 (MD3) and CFP-PARP15 (MD1) or to 16 °C for constructs CFP-SARS-CoV-2 and SARS-CoV-2 or to 18 °C for all other constructs and incubation continued overnight for about 20-22 hours. The cells were collected by centrifugation at 4,200×g for 15-30 min at 4 °C. The pellets were resuspended in respective lysis buffers (**Table S2**) and stored at -20 °C until purification.

#### Protein purification

##### IMAC

All constructs were initially purified by immobilized metal affinity chromatography (IMAC). The cells were thawed, supplemented with 0.1 mM Pefablock SC (Roche) and 20 µg/ml DNase I (Roche) and lysed by sonication. The lysate was centrifuged (16,000×g, 4 °C, 30 min), filtered and loaded onto an IMAC equilibrated with lysis buffer and charged with $Ni^{2+}$ or $Zn^{2+}$. The column was washed with lysis buffer and wash buffer and the protein was eluted with elution buffer. Detailed information with the column, buffers and volumes used for each construct can be found in **Table S2**.

##### Reverse IMAC

After purification by IMAC, a reverse IMAC step was performed for SARS-CoV-2 protein. The $His_6$-TrxT-tag was removed by digestion with TEV-protease (1:30 molar ratio) while the protein was dialyzed against 30 mM HEPES pH 7.5, 300mM NaCl, 10% glycerol, 0,5 mM TCEP (4 °C, 20 hours). The protein was thereafter loaded to a 5 ml HiTrap Chelating HP column charged with $Ni^{2+}$. The column was washed with 3 column volumes of lysis buffer. The flowthrough and wash fractions were collected for further purification by size-exclusion chromatography.

##### MBP affinity chromatography

After purification by IMAC, a final MBP affinity purification step was performed for MBP-tagged TNKS1-SAM-Catalytic construct. The IMAC eluate was loaded onto a 5 ml MBPTrap HP column equilibrated with MBPTrap loading buffer (50mM HEPES pH 7.5, 500 mM NaCl, 0.5mM TCEP). The column was washed with 3 column volumes of the MBPTrap loading buffer and eluted with the same buffer containing 10 mM maltose. The protein was aliquoted and flash frozen in liquid nitrogen and stored at -70 °C.

##### Size exclusion chromatography

With exception of Nluc-eAF1521 and MBP-TNKS1, all constructs were purified by a final size exclusion chromatography step. Size exclusion chromatography was carried out on a S75 16/600 size-exclusion chromatography column with the previously purified IMAC eluates or reverse IMAC fractions of the constructs. The buffer used was 20 mM HEPES pH 7.5, 300 mM NaCl, 0.5 mM TCEP for PtxS1, 20 mM HEPES pH 7.5, 250 mM NaCl, 1 mM TCEP, 5% glycerol for CFP-SARS-CoV, CFP-PARP9 (MD2) and CFP-PARP15 (MD2). For all other constructs, 30 mM HEPES (pH 7.5), 300 mM NaCl, 10% Glycerol, 0.5 mM TCEP was used. Purified fractions were combined, aliquoted and flash frozen in liquid nitrogen and stored at -70 °C.

**Table S2: IMAC purification information.** If only imidazole content is noted for the wash buffer or elution buffer, the other buffer components are identical to the respective lysis buffer used. We estimated the yield of protein after purification by IMAC per liter of culture. Wash volumes are shown in column volumes (CV).

| **Construct name** | IMAC column (charged with salt) | Lysis buffer (initial wash column volumes) | Wash buffer (imidazole, second wash column volumes) | Elution buffer | Approx. yield per liter culture |
|---|---|---|---|---|---|
| CFP-ALC1 | 5 ml HiTrap IMAC HP ($Ni^{2+}$) | 50 mM HEPES pH 7.5, 500 mM NaCl, 10 mM Imidazole, 10% glycerol, 0.5mM TCEP (10 CV) | none | 350 mM imidazole | 15 mg |
| CFP-APLF | 5 ml HiTrap Chelating HP ($Zn^{2+}$) | 50 mM HEPES pH 7.5, 500 mM NaCl, 10 mM Imidazole, 0.5 mM TCEP (4 CV) | 30 mM imidazole (4 CV) | 100 mM imidazole | >300 mg |
| CFP-ARH1<br>CFP-ARH2 | 5 ml HiTrap IMAC HP ($Ni^{2+}$) | 50 mM HEPES pH 7.5, 10 mM $MgCl_2$, 10 mM imidazole, 500 mM NaCl, 10% glycerol, 0.5mM TCEP (10 CV) | 20 mM imidazole (10 CV) &<br>30 mM imidazole (5 CV) | 100 mM imidazole | 100 mg |
| CFP-ARH3 | 5 ml HiTrap IMAC HP ($Ni^{2+}$) | 50 mM HEPES pH 7.5, 10 mM $MgCl_2$, 10 mM imidazole, 500 mM NaCl, 10% glycerol, 0.5mM TCEP<br>(10 CV) | 20 mM imidazole (10 CV) &<br>30 mM imidazole (2 CV) | 100 mM imidazole | >300 mg |
| CFP-CHIKV | 5 ml HiTrap Chelating HP ($Ni^{2+}$) | 50mM HEPES pH 7.5, 500 mM NaCl, 10 mM Imidazole, 10% glycerol, 0.5mM TCEP<br>(10 CV) | 20 mM imidazole (10 CV) &<br>30 mM imidazole (2 CV) | 100 mM imidazole | >300 mg |
| CFP-eAf1521 | 5 ml HiTrap Chelating HP ($Ni^{2+}$) | 50mM HEPES pH 7.5, 500 mM NaCl, 10 mM Imidazole, 0.5mM TCEP<br>(4 CV) | none | 100 mM imidazole | >300 mg |
| CFP-GDAP2 | 5 ml HiTrap IMAC HP ($Ni^{2+}$) | 50mM HEPES pH 7.5, 500 mM NaCl, 10 mM Imidazole, 10% glycerol, 0.5mM TCEP<br>(10 CV) | 20 mM imidazole (10 CV) &<br>30 mM imidazole (4 CV) | 100 mM imidazole | >300 mg |
| CFP-MacroH2A1.1<br>CFP-MacroH2A1.2<br>CFP-MacroH2A.2 | 5 ml HiTrap Chelating HP ($Ni^{2+}$) | 50 mM HEPES pH 7.5, 500 mM NaCl, 10 mM Imidazole, 0.5mM TCEP (9 CV) | 25 mM imidazole (9 CV) | 250 mM imidazole | >300 mg |
| CFP-MDO1 | 5 ml HiTrap Chelating HP ($Ni^{2+}$) | 50mM HEPES pH 7.5, 500 mM NaCl, 10 mM Imidazole, 10% glycerol, 0.5mM TCEP (10 CV) | 20 mM imidazole (10 CV) &<br>30 mM imidazole (10 CV) | 100 mM imidazole | 200 mg |
| CFP-MDO2 | 5 ml HiTrap IMAC HP ($Ni^{2+}$) | 50 mM HEPES pH 7.5, 500 mM NaCl, 10 mM Imidazole, 10% glycerol, 0.5 mM TCEP (4 CV) | none | 100 mM imidazole | >300 mg |
| CFP-MERS-CoV | 5 ml HiTrap Chelating HP ($Ni^{2+}$) | 50mM HEPES pH 7.5, 500 mM NaCl, 10 mM Imidazole, 10% glycerol, 0.5mM TCEP (10 CV) | 20 mM imidazole (4 CV) | 100 mM imidazole | 200 mg/L |
| CFP-PARG | 5 ml HiTrap Chelating HP ($Ni^{2+}$) | 50mM HEPES pH 7.5, 500 mM NaCl, 10 mM Imidazole, 10% glycerol, 0.5mM TCEP (10 CV) | 20 mM imidazole (10 CV) &<br>30 mM imidazole (10 CV) | 100 mM imidazole | 40 mg |

**Table S2: IMAC purification information. (continued)**

| **Construct name** | IMAC column (charged with salt) | Lysis buffer (initial wash column volumes) | Wash buffer (imidazole, second wash column volumes) | Elution buffer | Approx. yield per liter culture |
|---|---|---|---|---|---|
| CFP-PARP14 (MD1) | 5 ml HiTrap Chelating HP ($Ni^{2+}$) | 50mM HEPES pH 7.5, 500 mM NaCl, 10 mM imidazole, 10% glycerol, 0.5mM TCEP (10 CV) | 20 mM imidazole (10 CV) & 30 mM imidazole (10 CV) | 100 mM imidazole | 25 mg |
| CFP-PARP14 (MD2) | 5 ml HiTrap Chelating HP ($Ni^{2+}$) | 50mM HEPES pH 7.5, 500 mM NaCl, 10 mM imidazole, 10% glycerol, 0.5mM TCEP (10 CV) | 20 mM imidazole (10 CV) & 30 mM imidazole (10 CV) | 100 mM imidazole | 250 mg |
| CFP-PARP14 (MD3) | 5 ml HiTrap Chelating HP ($Ni^{2+}$) | 50mM HEPES pH 7.5, 500 mM NaCl, 10 mM imidazole, 10% glycerol, 0.5mM TCEP (10 CV) | 20 mM imidazole (10 CV) & 30 mM imidazole (5 CV) | 100 mM imidazole | 120 mg |
| CFP-PARP15 (MD1) | 5 ml HiTrap Chelating HP ($Ni^{2+}$) | 50mM HEPES pH 7.5, 500 mM NaCl, 10 mM imidazole, 0.5mM TCEP (5 CV) | 18 mM imidazole (5 CV) | 200 mM imidazole | 4 mg |
| CFP-PARP15 (MD2) | 5 ml HiTrap IMAC HP ($Ni^{2+}$) | 50mM HEPES pH 7.5, 500 mM NaCl, 10 mM imidazole, 10% glycerol, 0.5mM TCEP (6 CV) | 30mM HEPES pH 7.5, 500 mM NaCl, 25 mM Imidazole, 0.5mM TCEP, 10% glycerol (6 CV) | 30mM HEPES pH 7.5, 500 mM NaCl, 300 mM Imidazole, 0.5mM TCEP, 10% glycerol | >300 mg |
| CFP-PARP9 (MD1) | 5 ml HiTrap IMAC HP ($Ni^{2+}$) | 50 mM HEPES pH 7.5, 500 mM NaCl, 10 mM imidazole, 10% glycerol, 0.5 mM TCEP (7 CV) | 20 mM imidazole (10 CV) | 100 mM imidazole | 3 mg |
| CFP-PARP9 (MD2) | 5 ml HiTrap IMAC HP ($Ni^{2+}$) | 50 mM HEPES pH 7.5, 500 mM NaCl, 10 mM imidazole, 10% glycerol, 0.5 mM TCEP (6 CV) | 30mM HEPES pH 7.5, 500 mM NaCl, 25 mM Imidazole, 0.5mM TCEP, 10% glycerol (6 CV) | 30mM HEPES pH 7.5, 500 mM NaCl, 300 mM Imidazole, 0.5mM TCEP, 10% glycerol | 25 mg |
| CFP-RNF146 | 2 ml HisPur Ni-NTA resin | 50 mM HEPES pH 7.5, 500 mM NaCl, 10 mM imidazole, 10% glycerol, 0.5mM TCEP (25 CV) | 25 mM imidazole (25 CV) | 200 mM imidazole | >300 mg |
| CFP-SARS-CoV | 5 ml HiTrap Chelating HP ($Ni^{2+}$) | 50mM HEPES pH 7.5, 500 mM NaCl, 10 mM imidazole, 0.5mM TCEP, 10% glycerol (6 CV) | 30mM HEPES pH 7.5, 500 mM NaCl, 25 mM Imidazole, 0.5mM TCEP, 10% glycerol (6 CV) | 30mM HEPES pH 7.5, 500 mM NaCl, 300 mM Imidazole, 0.5mM TCEP, 10% glycerol (6 CV) | 200 mg |
| CFP-SARS-CoV-2 | 5 ml HiTrap Chelating HP ($Ni^{2+}$) | Buffer: 50mM HEPES pH 7.5, 500 mM NaCl, 10 mM imidazole, 10% glycerol, 0.5mM TCEP (15 CV) | 30 mM imidazole (7 CV) | Linear gradient over 10 CV (10-260 mM imidazole) | >300 mg |
| CFP-TARG1 | 5 ml HiTrap Chelating HP ($Ni^{2+}$) | 50mM HEPES pH 7.5, 500 mM NaCl, 10 mM imidazole, 10% glycerol, 0.5mM TCEP (10 CV) | 20 mM imidazole (10 CV) & 30 mM imidazole (2 CV) | 100 mM imidazole | 200 mg |
| CFP-XRCC1 | 2 ml HisPur Ni-NTA resin | 50 mM HEPES pH 7.5, 500 mM NaCl, 10 mM imidazole, 10% glycerol, 0.5mM TCEP (10 CV) | none | 350 mM imidazole | 10 mg |
| NanoLuc-ALC1 | 2 ml HisPur Ni-NTA resin | 50mM HEPES pH 7.5, 500 mM NaCl, 10 mM imidazole, 10% glycerol, 0.5mM TCEP (25 CV) | 25 mM imidazole (25 CV) | 200 mM imidazole | 100 mg |
| NanoLuc-eAf1521 | 5 ml HiTrap Chelating HP ($Ni^{2+}$) | 50mM HEPES pH 7.5, 500 mM NaCl, 10 mM imidazole, 0.5mM TCEP (4 CV) | none | 100 mM imidazole | 200 mg |

**Table S2: IMAC purification information. (continued)**

| **Construct name** | IMAC column (charged with salt) | Lysis buffer (initial wash column volumes) | Wash buffer (imidazole, second wash column volumes) | Elution buffer | Approx. yield per liter culture |
|---|---|---|---|---|---|
| NanoLuc-MDO2 | 2 ml HisPur Ni-NTA resin | 50mM HEPES pH 7.5, 500 mM NaCl, 10 mM imidazole, 10% glycerol, 0.5mM TCEP (25 CV) | 25 mM imidazole (25 CV) | 200 mM imidazole | 200 mg |
| PtxS1 | 1 ml HiTrap Chelating HP ($Ni^{2+}$) | 50mM HEPES pH 7.5, 500 mM NaCl, 10 mM imidazole, 10% Glycerol, 0.5mM TCEP (20 CV) | 25 mM imidazole (10 CV) | 350 mM imidazole | 10 mg |
| SARS-CoV-2 | 5 ml HiTrap Chelating HP ($Ni^{2+}$) | Buffer: 50mM HEPES pH 7.5, 500 mM NaCl, 10 mM imidazole, 10% glycerol, 0.5mM TCEP (30 CV) | 30 mM imidazole (4 CV) | 300 mM imidazole | 50 mg |
| TNKS 1 SAM-Catalytic (E1050K) | 5 ml HiTrap IMAC HP ($Ni^{2+}$) | 50mM HEPES pH 7.5, 500 mM NaCl, 20 mM imidazole, 10% glycerol,0.5mM TCEP (10 CV) | 50 mM imidazole (5 CV) | 500 mM imidazole | 5 mg |
| TNKS1 SAM-Catalytic | 5 ml HiTrap Chelating HP ($Ni^{2+}$) | 50mM HEPES pH 7.5, 500 mM NaCl, 15 mM imidazole, 0.5mM TCEP (10 CV) | 35 mM imidazole (8 CV) | 300 mM imidazole | 30 mg |
| TNKS1 SAM-Catalytic (Y1073A) | 5 ml HiTrap IMAC HP ($Ni^{2+}$) | 40mM HEPES pH 7.5, 500 mM NaCl, 20 mM imidazole, 10% glycerol,0.5mM TCEP (8 CV) | 50 mM imidazole (10 CV) | 500 mM imidazole | 5 mg |
| YFP-GAP | 5 ml HiTrap IMAC HP ($Ni^{2+}$) | 50 mM HEPES pH 7.5, 500 mM NaCl, 10 mM imidazole, 10% glycerol,0.5 mM TCEP (10 CV) | 10 mM imidazole (5 CV) | 350 mM imidazole | >300 mg |
| YFP-GAP(C→A) | 5 ml HiTrap IMAC HP ($Ni^{2+}$) | 50 mM HEPES pH 7.5, 500 mM NaCl, 10 mM imidazole, 10% glycerol,0.5 mM TCEP (10 CV) | 10 mM imidazole (5 CV) | 350 mM imidazole | >300 mg |
| YFP-G$\alpha_i$ (full length) | 5 ml HiTrap IMAC HP ($Ni^{2+}$) | 50 mM HEPES pH 7.5, 500 mM NaCl, 10 mM imidazole, 10% glycerol, 0.5 mM TCEP (3 CV) | 20 mM imidazole (10 CV) | 300 mM imidazole | 10 mg |

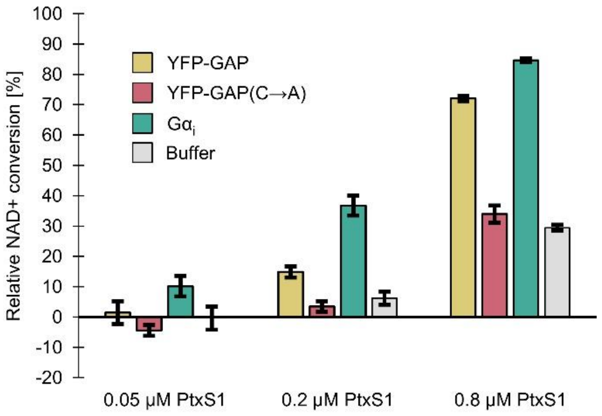


**Figure S1: NAD$^+$ consumption by PtxS1 with different Gα$_i$ constructs.** 0.05 µM, 0.2 µM or 0.8 µM PtxS1 were mixed with 30 µM NAD$^+$ and 20 µM of YFP-GAP, YFP-GAP(C→A), full length Gα$_i$ or buffer. Samples were incubated for 1 hour at room temperature. Data shown are mean ± standard deviation with number of replicates n = 4.

The NAD consumption assay for PtxS1 was performed as previously described (Ashok et al., 2020). All incubation steps were performed at room temperature. Briefly, reactions were mixed and 10 µl per well transferred to a black polypropylene low-volume 384-well plate (FisherBrand) and incubated for 1 h. Samples containing no PtxS1 were used to calculate the relative conversion. Unreacted NAD$^+$ was converted to a fluorescent product by addition of 4 µl 2 M KOH and 4 µl 20%(v/v) acetophenone in ethanol. After incubation for 10 minutes, 18 µl formic acid per well was added and reactions were incubated for 30 minutes. Fluorescence was measure using Tecan M1000 Pro multimode plate reader. The samples were excited at 372 nm wavelength and emission at 444 nm wavelength was measured.

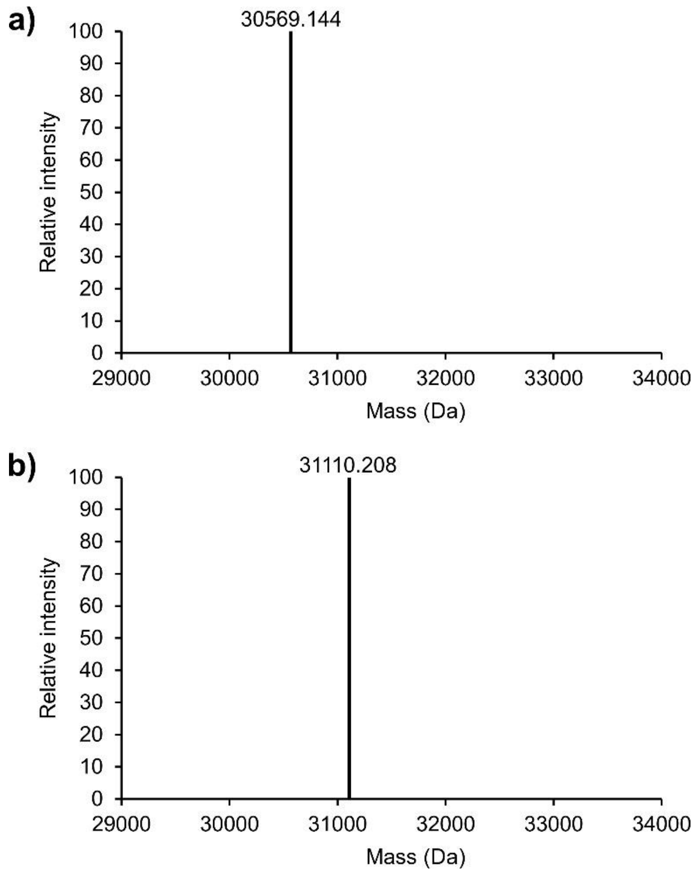


**Figure S2: Analysis of YFP-GAP ADP-ribosylation by mass spectrometry.** The deconvoluted monoisotopic mass spectra are shown for YFP-GAP (a) before and (b) after ADP-ribosylation with PtxS1 and subsequent purification. The mass difference is 541.064 Da and corresponds to the theoretical monoisotopic mass of a single ADP-ribosyl group (541.061 Da).

50 µM of YFP-GAP or YFP-GAP(MAR) were mixed with 0.1% trifluoracetic acid. The molecular weights of purified protein samples were measured by electrospray ionization mass spectrometry combined with liquid chromatography (LC-ESI-MS) using a Q Exactive Plus Mass Spectrometer.

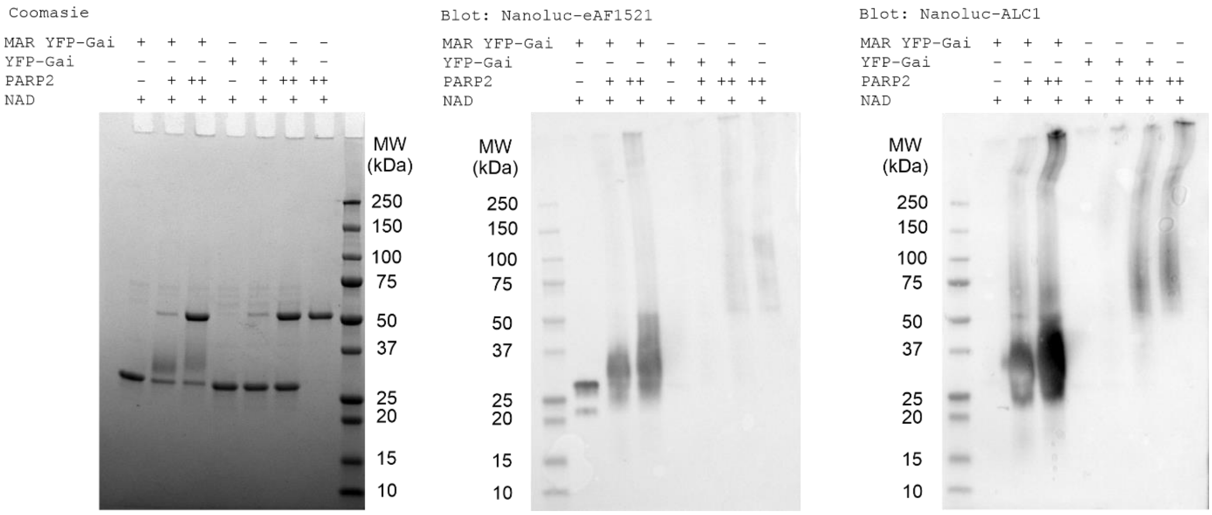


**Figure S3: PARP2 can extend modify YFP-GAP-MARylated to produce a PARylated form.** Full gel and blots of those shown in figure 2B.

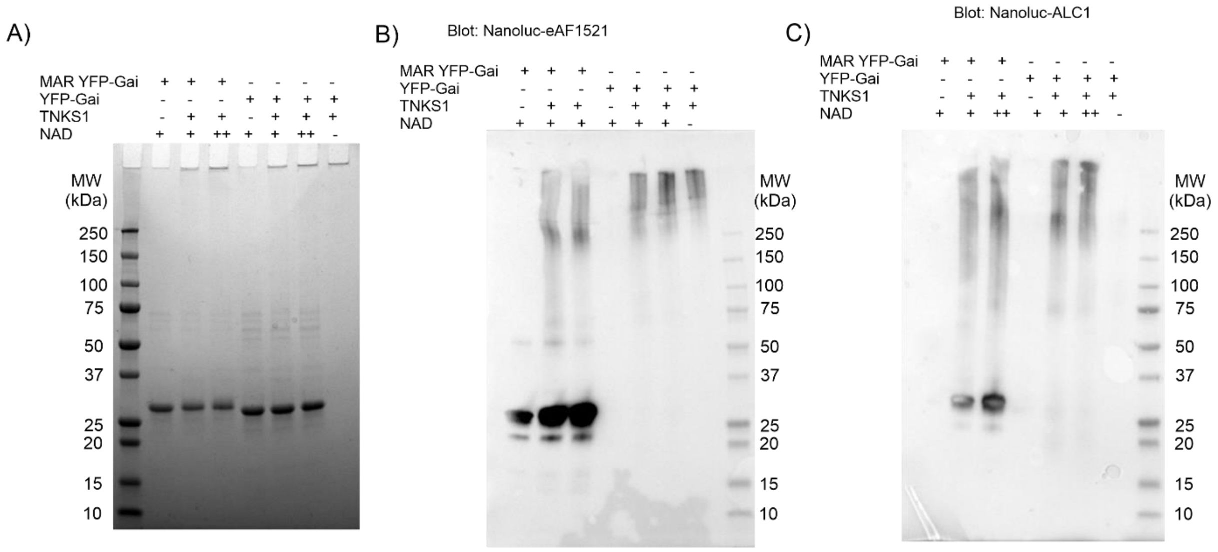


**Figure S4: PARylation of YFP-GAP by TNKS1.** 200nM TNKS1 (SAM-Catalytic domain dimer) was mixed with YFP-GAP or YFP-GAP MARylated and incubated overnight at room temperature in the presence of 1 mM (+) or 10 mM (++) $NAD^+$. Samples were analysed by SDS-PAGE and coomassie staining (A), as well as with western blot using nanoLuc-eAF1520 (B) or nanoLuc-ALC1 (C).

Dimeric TNKS1 was formed by incubating 2 independently purified samples consisting of TNKS1 SAM-Catalytic domain containing mutations E1050K and Y1073SA in SAM domain. Incubation of YFP-GAP is MARylated with TNKS1 dimer results in a change in electrophoretic mobility (A). The PARylation of YFP-GAP requires the protein to be previously MARylated suggesting that only one PAR chain is added to the YFP-GAP in extending the MAR modification. Since ALC1 detects only PAR chains, the blot in (C) indicates that the modification formed corresponds to a PAR polymer.

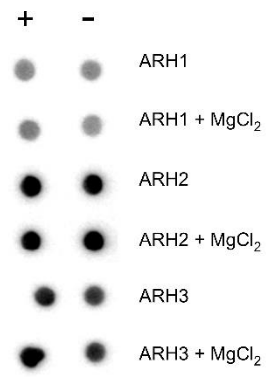


**Figure S5: Hydrolysis test of cysteine-ADP-ribose with ARH family members.** 10 µM of YFP-GAP(MAR) were incubated in presence (+) or absence (-) of 1 µM CFP-fused ARH constructs. The reactions were prepared in the presence of absence of 5 mM $MgCl_2$, incubated for 24 h at room temperature and thereafter blotted on nitrocellulose membranes. The membranes were washed and the protein-bound ADP-ribosyl-groups detected using Nluc-eAf1521.

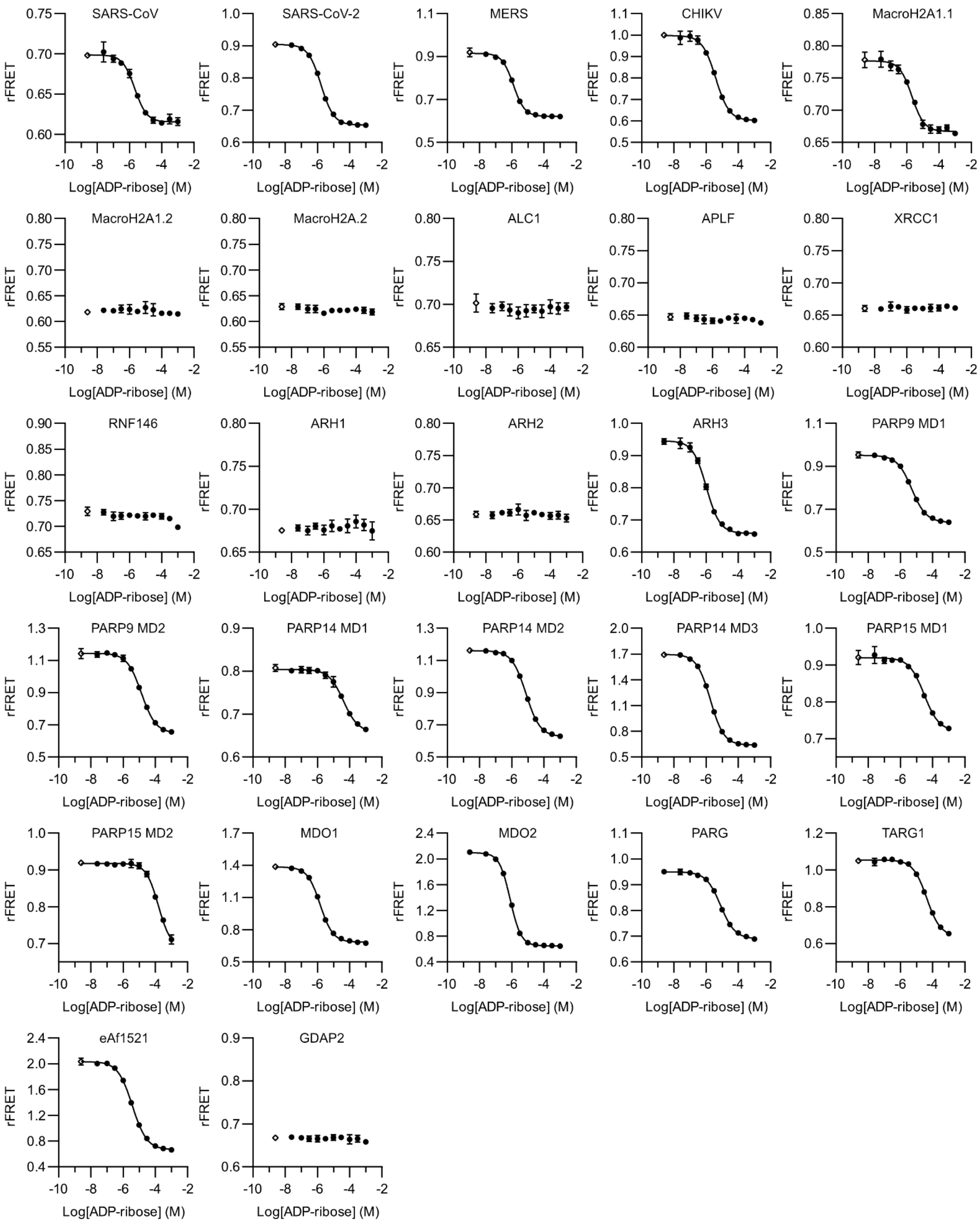


**Figure S6: ADP-ribose dose-response curves for CFP-fusion constructs and MARylated YFP-GAP.** 1 µM of CFP-fused constructs were mixed with 5 µM of YFP-GAP(MAR) and increasing concentrations of ADP-ribose. The ratiometric FRET signals (rFRET) were determined. Data shown are mean ± standard deviation with number of replicates n = 4. The controls containing no ADP-ribose were set one logarithmic unit below the lowest concentration.

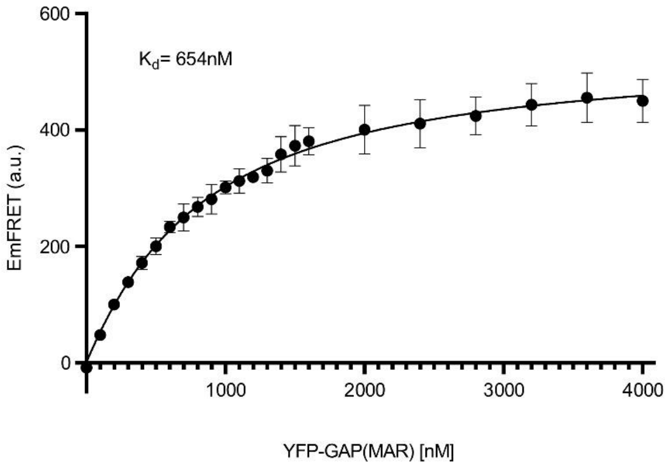


**Figure S7: Determination of binding affinity for CFP-MDO2 and YFP-GAP(MAR) by FRET.** 290 nM CFP-MDO2 were mixed with increasing concentrations of YFP-GAP(MAR). Determination of the binding affinity was done as previously described (Sowa et al., 2020).

a)

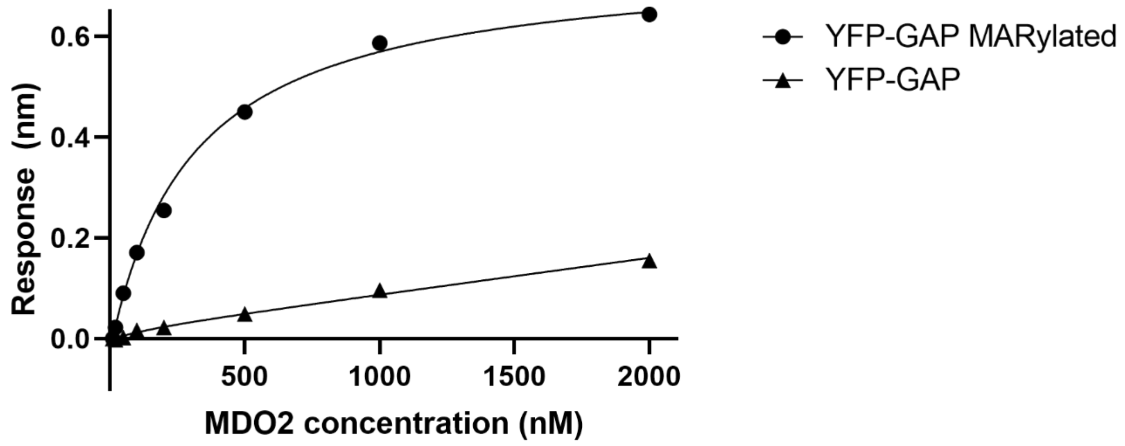


b)

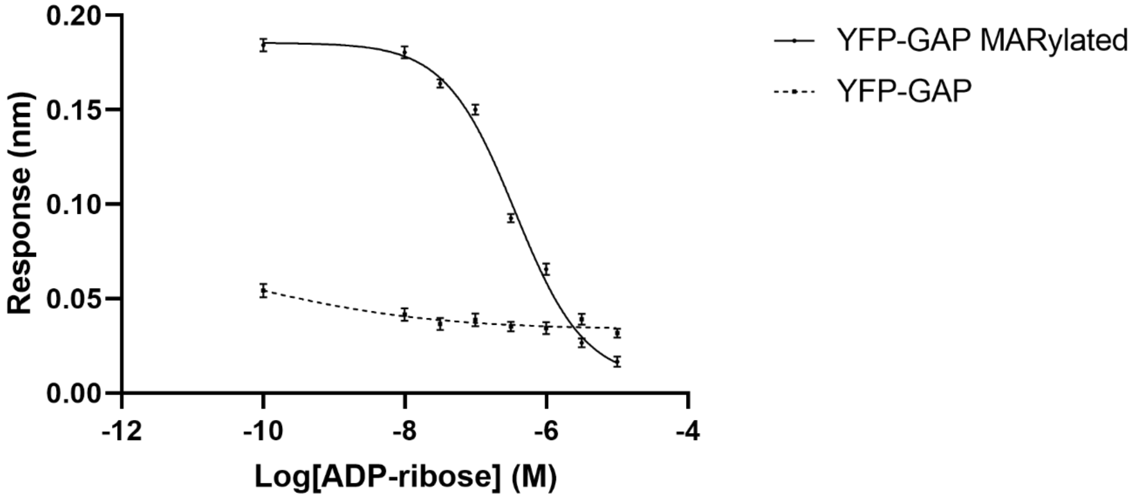


**Figure S8: Use of BLI to measure MDO2 to YFP-GAP-MARylated interaction.** A) Affinity of MDO2 for YFP-GAP-MARylated loaded sensors. Steady state signal is plotted against MDO2 concentration in solution. $K_D$ 500 ±120 nM for YFP-GAP-MARylated loaded sensor was determined with single exponential fit to 3 independent experiments. Individual values represent the average ± SD response during last 20 seconds of the association step. B) $IC_{50}$ determination of ADP-ribose with MDO2. MDO2 concentration was kept constant at 100 nM and ADP-ribose concentration was varied between 10 µM to 10 nM. $IC_{50}$ determined for YFP-GAP MARylated is 440 nM($pIC_{50}$=6.36 ± 0.08) from 3 independent experiments. Individual values represent the average ± SD response during last 20 seconds of the association step.
